# Robust organ size in Arabidopsis is primarily governed by cell growth rather than cell division patterns

**DOI:** 10.1101/2023.11.11.566685

**Authors:** Isabella Burda, Chun-Biu Li, Frances K. Clark, Adrienne H. K. Roeder

**Affiliations:** Genetics, Genomics, and Development Graduate Program, Cornell University, Ithaca, NY 14850, USA; Weill Institute for Cell and Molecular Biology Cornell University, Ithaca, NY, 14850, USA; School of Integrative Plant Science, Section of Plant Biology, Cornell University, Ithaca, NY, 14850, USA; Department of Mathematics, Stockholm University, Stockholm 10691, Sweden

**Keywords:** *Arabidopsis thaliana*, sepal, spatiotemporal averaging, *FTSH4*, endoreduplication, LGO, morphogenesis, giant cells

## Abstract

Organ sizes and shapes are highly reproducible, or robust, within a species and individuals. *Arabidopsis thaliana* sepals, which are the leaf-like organs that enclose flower buds, have consistent size and shape, which indicates robust development. Counterintuitively, variability in cell growth rate over time and between cells facilitates robust development because cumulative cell growth averages to a uniform rate. Here we investigate how sepal morphogenesis is robust to changes in cell division but not robust to changes in cell growth variability. We live image and quantitatively compare the development of sepals with increased or decreased cell division rate (*lgo* mutant and *LGO* overexpression, respectively), a mutant with altered cell growth variability (*ftsh4*), and double mutants combining these. We find that robustness is preserved when cell division rate changes because there is no change in the spatial pattern of growth. Meanwhile when robustness is lost in *ftsh4* mutants, cell growth accumulates unevenly, and cells have disorganized growth directions. Thus, we demonstrate *in vivo* that both cell growth rate and direction average in robust development, preserving robustness despite changes in cell division.

**Summary statement:** Robust sepal development is preserved despite changes in cell division rate and is characterized by spatiotemporal averaging of heterogeneity in cell growth rate and direction.

## Introduction

Many aspects of development are robust, meaning that they have reproducible outcomes despite internal or environmental noise. Many organs have robust size and shape, which is important for proper function (Hong et al, 2018, Boulan and Leopold, 2021). *Arabidopsis thaliana* (hereafter Arabidopsis) sepals, which are the leaf-like organs that enclose flower buds, have uniform size and shape which indicates that development of size and shape is robust. (Hong et al, 2016; Zhu et al, 2020).

Organ size and cell size can be uncoupled during development, which is called compensation. It has often been observed that changing cell size does not proportionally change organ size. Instead, an organ with less cells will have larger cell sizes, and an organ with more cells will have smaller cells. Many mutants that exhibit compensation in leaf development have fewer cells per leaf, but those cells are proportionally larger in size (Ferjani et al, 2007; Horiguchi and Tsukaya, 2011). The same trade-off between cell size and cell number occurs in Drosophila wing development (Neufeld et al, 1998). Compensation is linked to a long-standing debate of whether cells or organs are considered the basic unit of plants (Kaplan and Hagemann, 1991; Kaplan 1992) and indicates that control of organ size and shape is complex.

Compensation has been observed in the Arabidopsis sepal when there is a change in the rate of endoreduplication (Roeder et al, 2010; Robinson et al, 2018). Endoreduplication is an alternative to mitosis in which DNA replication occurs without cell division producing a cell with increased ploidy (Trass et al, 1998; Veydler et al, 2011). Endoreduplication is necessary for the differentiation of specialized epidermal cell types such as trichomes and giant cells (Churchman et al, 2006; Van Leene et al, 2010; Kumar et al, 2015, Roeder et al, 2010; Robinson et al, 2018). The sepal epidermis has endoreduplicated giant cells which are interspersed among smaller epidermal cells (Roeder et al, 2010). Overexpression of the cyclin-dependent kinase inhibitor LOSS OF GIANT CELLS FROM ORGANS (LGO), also called SMR1, results in an increased number of giant cells that have undergone endoreduplication whereas *lgo-2* mutants have few to no giant cells (Roeder et al, 2010; Schwarz and Roeder, 2016; Kumar et al, 2015). However, *LGO* overexpression (*pATML1::LGO*; hereafter *LGOoe*) sepals have the same area as wild-type sepals, and *lgo-2* mutant sepals are only slightly larger in area than wild type (Robinson et al, 2018). This occurs because *LGOoe* sepals have fewer cells and *lgo-2* sepals have more cells (Roeder et al, 2010; Robinson et al, 2018). Thus, compensation occurs to preserve final organ size. However, sepal size and shape also need to be uniform throughout their development to stay closed as the flower grows. It is not well understood how the sepal compensates for the change in cell division during development.

Mutants with variable organ size and shape have been used to elucidate mechanisms generating reproducible organ size and shape. For example, sepal size and shape are variable in the *drmy1* mutant due to altered timing of the initiation of primordia from the floral meristem. Disrupted patterning of cytokinin and auxin underlie the abnormal timing of primordia initiation (Zhu et al, 2020; Kong et al, 2023). Thus, nearly synchronous initiation of sepals promotes robust sepal size. Variable sepal size and shape also occurs in *vip3*, due to noisy transcription (Trinh et al, 2023). In addition, the mitochondrial protease mutant *ftsh4-5* has variable sepal size and shape which results from elevated levels of reactive oxygen species (Hong et al, 2016). FTSH4 (Filamentous temperature sensitive H 4) is an iAAA-protease located in the inner mitochondrial membrane. Both its protease activity and chaperone activity have roles in eliminating aggregated and carbonylated proteins in the mitochondria (Maziak et al, 2021) and *ftsh4* mutants also have abnormal mitochondrial morphology (Gibala et al, 2009). Besides the variable organ size and shape, a variety of developmental phenotypes have been reported in *ftsh4* mutants including delayed bolting (Gibala et al, 2009; Dolzblasz et al 2016), delayed germination, and lower leaf production (Gibala et al, 2009) in *ftsh4-1* and *ftsh4-2* and dwarfism and axillary branching (Zhang et al, 2014) in *ftsh4-4*. Other developmental phenotypes occur when *ftsh4* mutants are grown under heat stress conditions, such as shorter stems and lack of siliques (Dolzblasz et al, 2016). Short day conditions also cause developmental phenotypes in *ftsh4* mutants, such as serrated leaves with abnormal patterning of palisade cells and spongy mesophyll (Gibala et al, 2009). The phenotypes are ameliorated by decreasing reactive oxygen species (Zhang et al, 2014; Hong et al, 2016). Further, signs of oxidative stress increase with age in *ftsh4* (Gibala et al, 2009; Dolzblasz et al, 2016), and the abnormal morphology of *ftsh4* leaves is also associated with increased levels of reactive oxygen species as the plant ages (Gibala et al, 2009). These findings indicate that the developmental phenotypes, including loss of robust sepal development, are linked to the loss of function of *FTSH4* and consequent increases in reactive oxygen species (Gibala et al, 2009; Hong et al, 2016).

Counterintuitively, robust development is linked to heterogeneity in growth rates. Nearby epidermal cells can have up to four-fold difference in growth rate (Elsner et al, 2012). Since cell walls prevent plant cells from moving relative to each other, heterogeneity is generated at a subcellular scale, with portions of the cell wall within a cell growing at different rates (Elsner et al, 2012). During sepal development, epidermal cell growth rates vary both temporally (over the course of development) and spatially (between cells at a given developmental time) (Tauriello et al, 2015; Hong et al, 2016; Le Gloanec et al, 2022). Variability also results from differentiation of different epidermal cell types (Le Gloanec et al, 2022). The averaging of spatial and temporal variability into even growth is termed spatiotemporal averaging (Hong et al, 2016). Interestingly, there is decreased heterogeneity in cell growth rates during *ftsh4-5* sepal development (Hong et al, 2016), suggesting that heterogeneity facilitates robust development (Hong et al, 2016). Modeling indicates that robust development occurs because heterogeneity that is spatially and temporally random averages over time and ensures even growth throughout the organ (Hong et al, 2016).

Heterogeneity in growth rates has also been found to be important in other developmental contexts. The microtubule severing protein mutant *katanin* has less heterogeneity in growth rates and has abnormal morphology of organ primordia (Uyttewaal et al, 2012). Modeling suggests that the ability for microtubules to reorient in response to tension, and thus resist growth in the direction of tension, increases heterogeneity in growth rates. The ability to amplify heterogeneity likely allows primordia emergence from the meristem, because the primordia grows faster than the boundary region (Uyttewaal et al, 2012). Heterogeneous cell growth rates also occur in response to differentiation of trichomes and serve to preserve organ shape despite variation in trichome number (Hervieux et al, 2017). The initial fast growth and then subsequent slow growth of trichomes causes nearby cells to restrict their growth rates, thus acting as a buffer and preventing organ shape change (Hervieux et al, 2017, Le Gloanec et al, 2022). Thus, heterogeneity could be a response to noise and cell type differentiation during development and facilitate robustness of organ size and shape development.

Here, we test the role of cell division in robustness development by increasing and decreasing cell division rate in the wild type and *ftsh4-5* background. We use *LGO* expression level to modulate cell division rate and *ftsh4-5* to alter robustness. Then we perform time lapse imaging to understand how growth is affected in the developing sepal. We find that the spatial pattern of cell growth and cell growth direction are the same in WT, overexpression of *LGO*, and a *lgo* null mutant despite changes in cell size. The preserved pattern of cell growth and cell growth direction explains how sepal morphogenesis is robust to changes in cell division. In contrast, the spatial pattern of cell growth and cell growth direction are altered by *ftsh4-5* and double mutants. Further, in the wild-type background, the variability of growth averages over time to produce even growth and organized growth direction despite changes in division rate. However, in the *ftsh4-5* background, cell growth accumulates unevenly over time and cells grow in disorganized directions. The defects in *ftsh4-5* cell growth are not ameliorated or accentuated by changes in cell division rate. Together, our results suggest that robust sepal development is driven by localization of growth rather than cell division and shows in vivo that heterogeneity in growth averages to produce uniform growth.

## Results

### Robustness of sepal size and shape is not affected by cell division

To test whether cell division affects robustness of sepal size and shape, we used the loss of function *lgo-2* allele, which increases cell division, and the gain of function *pATML1::LGO* transgenic plants (hereafter referred to as *LGOoe*), which decreases cell division in the sepal epidermis. Wild type (WT), *lgo-2*, and *LGOoe* have sepals that appear uniform in size and shape (Figure 1A-C). Double mutants were made with *ftsh4-5,* which has sepals of variable size and shape (Hong et al, 2016)(Figure 1D). In the mature flower, both *lgo-2 ftsh4-5* and *LGOoe ftsh4-5* have variable sepal size and shape similar to *ftsh4* single mutant, indicating that the *ftsh4-5* morphology remains when cell division rate changes (Figure 1E-F). During flower development, four sepals enclose each flower bud, and the sepals must have robust size and shape to maintain closure of the bud. WT and *lgo-2* buds are closed (Figure 1G-H) whereas *LGOoe* buds often have small gaps between adjacent sepals (Figure 1I) due to decreased sepal width which prevents the sepals from fully wrapping around the flower and was previously reported (Roeder et al, 2012). In contrast, *ftsh4-5*, *lgo-2 ftsh4-5*, and *LGOoe ftsh4-5* often have large gaps between sepals, particularly when buds have sepals with large differences in size or shape (Figure 1J-L). Therefore, the phenotypes of WT, *LGOoe*, and *lgo-2* are indicative of robust sepal development whereas the phenotypes of *ftsh4-5*, *lgo-2 ftsh4-5*, and *LGOoe ftsh4-5* are indicative of a loss of robustness.

**Figure 1:**
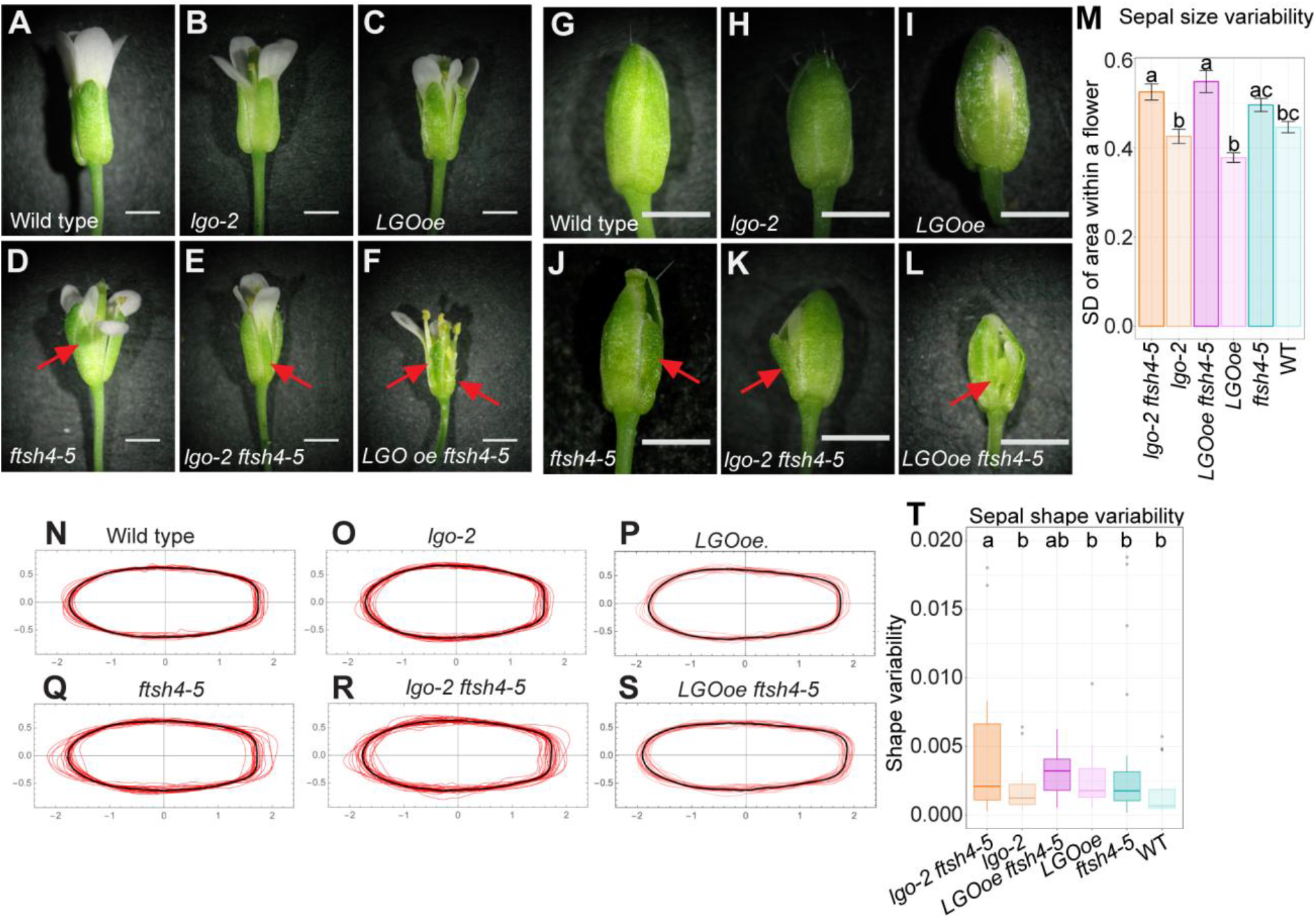
Organ size and shape is robust to changes in *LGO* expression, but not in *ftsh4-5* and double mutants. (A-F) Stage 15 mature flowers and (G-L) stage 12 flower buds of WT (A, G), *lgo-2* mutant (B, H), *LGOoe* (C, I; *pML1::LGO*), *ftsh4-5* mutant (D, J), *lgo-2 ftsh4-5* double mutant (E, K), and *LGOoe ftsh4-5* (F, L). Scale bars are 1 mm. Red arrows point to abnormally shaped sepals. (M) Bar graphs of standard deviation of sepal area within one flower. n=104 (WT), 100 (*lgo-2*), 104 (*LGOoe*), 108 (*ftsh4-5*), 116 (*lgo-2 ftsh4-5*), 108 (*LGOoe ftsh4*) sepals. Error bars show the standard error of the mean. Letters mark the groups that are not significantly different. (N-S) Contours of mature outer (abaxial) sepals normalized by size (red lines) and the average sepal shape (black line). n=18 (WT), 23 (*lgo-2*), 22 (*LGOoe*), 24 (*ftsh4-5*), 27 (*lgo-2 ftsh4-5*), 22 (*LGOoe ftsh4*) sepals. (T) Boxplots of variability of abaxial sepal shape (S_2 as described in Hong et al 2016). n=77 (WT), 66 (*lgo-2*), 78 (*LGOoe*), 85 (*ftsh4-5*), 96 (*lgo-2 ftsh4-5*), 74 (*LGOoe ftsh4*) The boxes extend from the lower to upper quartile values of the data with the midline indicating the median and the whiskers extend past 1.5 × interquartile range. Small dots for each box indicate outliers. Letters mark the groups that are not significantly different. Contours for inner (adaxial) and lateral sepals are available in Supplemental Figure S1.

To quantify robustness, all four sepals were dissected from mature flowers (n=25), photographed, and then contours outlining the sepal shapes were segmented from the photographs. To quantify the variability in size within a flower, standard deviation of sepal area within one flower was calculated. WT, *lgo-2,* and *LGOoe* have similar levels of variability in sepal size, and have less variability in sepal size compared to *ftsh4-5, lgo-2 ftsh4-5*, and *LGOoe ftsh4-5* respectively (Figure 1M) (the difference between WT and *ftsh4-5* shows the same trend as the other but does not reach statistical significance). To quantify variability in shape, contours were normalized by size. WT, *lgo-2,* and *LGOoe* (Figure 1N-P, S1A-C) have similar levels of variability around the average sepal shape, and have less variability compared to *ftsh4-5, lgo-2 ftsh4-5*, and *LGOoe ftsh4-5* respectively (Figure 1Q-T, SD-F) (the difference between *lgo-2* and *lgo-2 ftsh4-5* is statistically significant and the others how the same trend but do not reach statistical significance). Low variability in sepal size and shape explains the closure of the flower bud in WT, *lgo-2* as well as the mostly closed flower buds in *LGOoe.* Increased variability in sepal size and shape explains the opened flower buds in *ftsh4-5*, *lgo-2 ftsh4-5*, and *LGOoe ftsh4-5*. Our results show that uniformity of sepal size and shape within a flower is preserved when cell division rate is increased or decreased. Similarly, the *ftsh4-5* variability of sepal size and shape is unaffected by cell division rate.

### *LGO* expression changes cell division rate in WT and *ftsh4-5* backgrounds during development

To determine how sepal shape robustness is preserved despite extreme changes in cell division rate, we time lapse imaged living sepals during development. Sepals from each genotype were imaged every 24 hours for 6 days (n=3). Abaxial sepals were used because they face outwards, making them the most accessible for imaging. Flowers at stage 5 of development, when the sepals are about to enclose the floral meristem (Smyth, 1990), were chosen for the start of time lapse imaging. This time series captures earlier stages of development than had been imaged previously (Hong et al, 2016), and it spans the time during which *ftsh4-5* phenotype first becomes visible. MorphoGraphX (Barbier de Reuille et al 2015; Strauss et al, 2022) was used for segmentation of cell and lineage tracking in two and a half dimensions on the curved surface of the sepal.

To measure the extent to which genotype changed cell division rate during the time-lapse, the number of daughter cells per lineage over the 6-day time series was calculated. WT has a combination of giant cells that never divide that are interspersed with dividing lineages with smaller cells (Fig 2A, S2A). *lgo-2* has more daughter cells per lineage, and few to no cells that never divide (Fig 2B, S2B). *LGOoe* has fewer daughter cell per lineage and many non-dividing giant cells (Figure 2C, S2C). Thus, the expression level of *LGO* successfully modulates cell division rate. *ftsh4-5* (Figure 2D, S2D) has slightly fewer daughter cells per lineage than WT, and a similar amount of nondividing lineages. *lgo-2 ftsh4-5* has more daughter cells per lineage than *ftsh4-5*, and few to no non-dividing cells (Figure 2E, S2E). *LGOoe ftsh4-5* has fewer daughter cells per lineage than *ftsh4-5,* and mostly non-dividing giant cells (Figure 2F, S2F). Our results demonstrate that LGO expression level successfully modulates division rate in the *ftsh4-5* background as well as WT (Fig 2G). The count of nondividing cells in each genotype follows the same trend as the division rate but does not reach significance (Fig 2H). We conclude the phenotypes of the double mutants are additive, indicating that *LGO* and *FTSH4* function in separate pathways. Our results confirm that these genotypes can be used to test how cell division affects robustness in young developing sepals.

**Figure 2:**
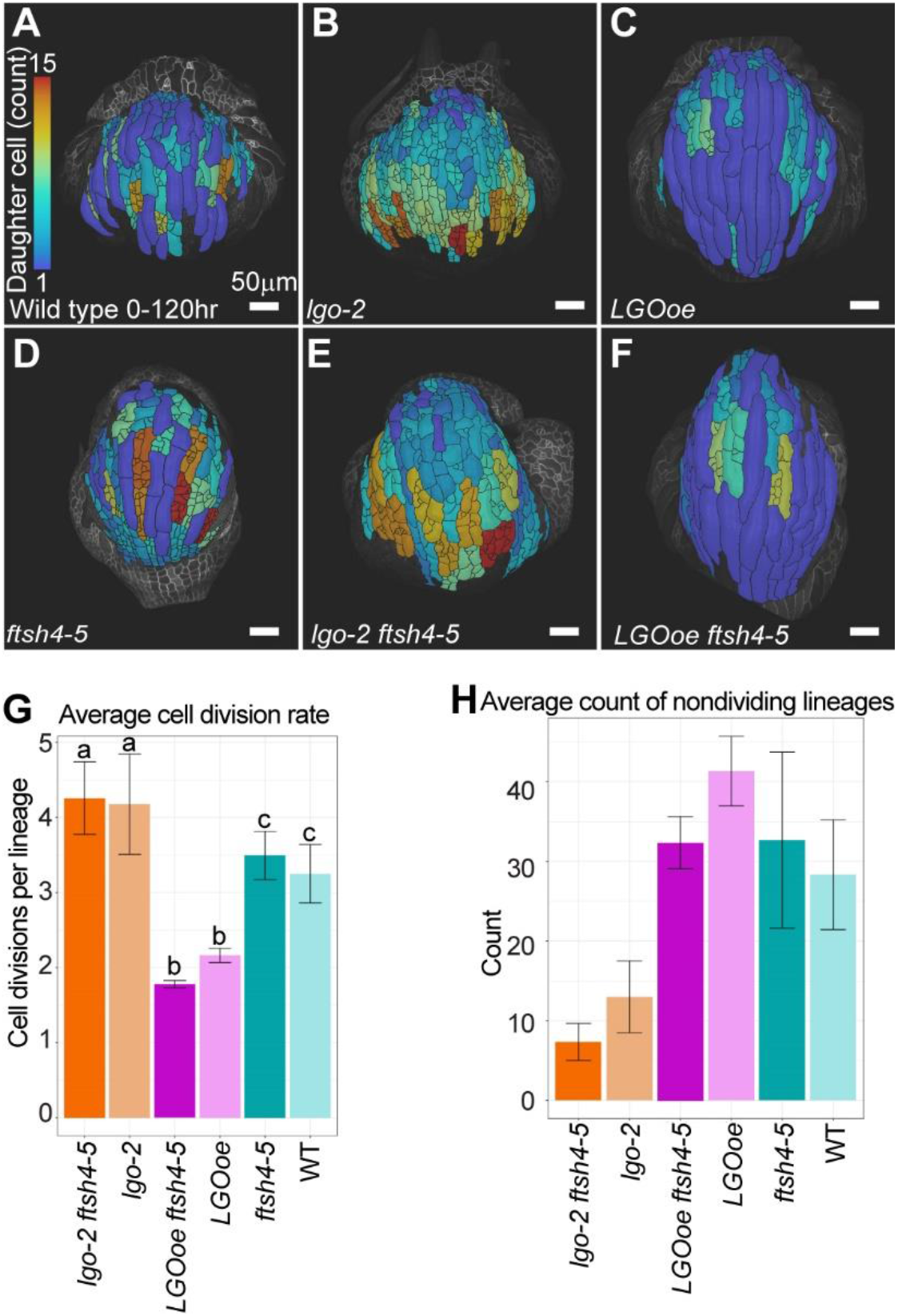
Cell division rate is decreased by *LGO overexpression* (*LGOoe*) and *LGOoe ftsh4* and is increased in *lgo-2* and *lgo-2 ftsh4-5*. (A-F) Heat maps of number of daughter cells per lineage using lineage tracking from 0-hour time point to 120-hour time point that are projected onto the 120-hour time point for WT (A), *lgo-2* (B), *LGOoe* (C), *ftsh4-5* (D), *lgo-2 ftsh4-4* (E), and *LGOoe* (F). The heat map scale is 1 to 15 daughter cells, where 1 indicates that no divisions have taken place because one cell gave rise to one cell at the final time point. The scale bar is 50µm. Representative images from n=3 biological replicates. Additional replicates are available in Supplemental Figure S2A-F. (G) Average number of daughter cells per lineage over the 120hr time lapse imaging. n=3. Error bars are the standard error of the mean. Letters mark the groups that are not significantly different. (H) Average number of cells that do not divide over the 120hr time lapse imaging. n=3. Error bars are the standard error of the mean.

### Cell division rate progressively changes cell size during development

To determine how changes in cell division affect cell size during development, cell area was measured at each time point throughout the course of time lapse imaging. At the start of the time lapse imaging, cell areas of WT, *lgo-2* and *LGOoe* are relatively homogenous and similar between genotypes. The mean cell size of *lgo-2* is the smallest at 99.4 µm^2^, the mean cell size of *LGOoe* is the largest at 157 µm^2^, and the mean cell size of WT is 106 µm^2^ (Figure 3A,D, Figure S3A-F). At the 48 and 72 hr time points, WT has a larger range of cell areas due to the differentiation of giant cells interspersed among smaller cells (Fig 3A,D S3A-B). The WT cell size distribution continues to widen in later time points (Fig 3A, S3M). The largest cells are differentiating into giant cells, which continue to endoreduplicate and grow in area (Roeder et al, 2010). Cell areas of *lgo-2* remain smaller and more homogenous than WT (Fig 3B,D FigS3C-D, M) and cell areas of LGOoe, get progressively larger (Fig3C-D Fig3SE-F,M). At the final time point, the mean cell size of *lgo-2* is the smallest at 221 µm^2^, the mean cell size of *LGOoe* is the largest at 797 µm^2^, and the mean cell size of WT is 342 µm^2^. *ftsh4-5* (Fig 3D-E, Fig S2G-H), *lgo-2 ftsh4-5* (Fig 3D,F, Fig S3 I-J), and *LGOoe ftsh4-5* (Fig 3D-G, Fig S3 K-M) mirror the cell area distributions of WT, *lgo-2* and *LGOoe* respectively. Therefore, the trade-off between cell size and cell division rate becomes pronounced during these developmental stages. Multidimensional scaling, which represents the differences in distributions as 2D distances, was used to analyze the cell areas in each time point and genotype. This reveals that cell areas of all genotypes cluster together at the 0 hr and 24 hr time points. *LGOoe* and *LGOoe ftsh4-5* no longer cluster with the other genotypes starting at the 48 hr time point. Then at the 72, 96, and 120 hr time points, WT and *ftsh4-5* cluster while *lgo-2* and *lgo-2 ftsh4-5* form another cluster (Figure S4). In summary, genotypes with different cell division rates have similar distributions of cell areas at the beginning of the time lapse imaging and become progressively different over time. The distribution of cell areas is not affected by *ftsh4-5.* Therefore, over the course of the time lapse, there is divergence in cell size distributions that is dependent on *LGO* expression level but not on *ftsh4-5*.

**Figure 3:**
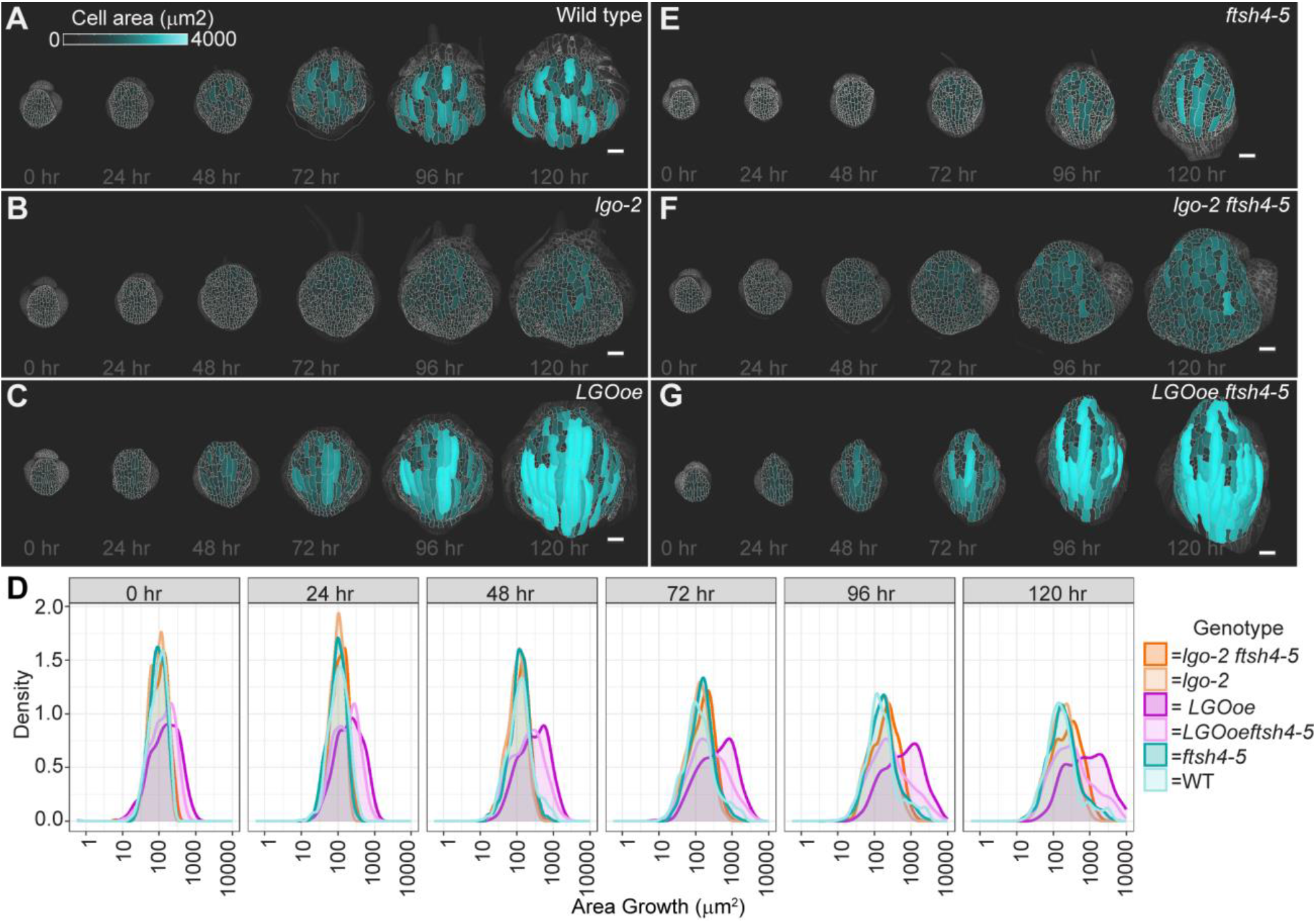
Cell sizes remain smaller in *lgo-2*, and *lgo-2 ftsh4-5,* and become progressively larger in *LGOoe* and *LGOoe ftsh4*. (A-G) Heat maps of cell area at each image time point for WT (A), *lgo-2* (B), *LGOoe* (C), *ftsh4-5* (D), *lgo-2 ftsh4-5* (E), and *LGOoe ftsh4-5* (F). The heat map scale is 0 to 4000µm^2^ and the scale bar is 50µm. Representative images from n=3 biological replicates. Additional replicates are available in Supplemental Figure S3. (D) Distribution of cell areas at each time point. Statistical analysis (multidimensional scaling) available in Supplemental Figure S4.

### Changing division rate does not change spatial localization of divisions within the developing sepal

To further characterize differences in cell division based on *LGO* expression and *ftsh4-5*, we examined the spatial localization of cell division over 24-hour time intervals. Cell divisions are represented as the change in number of cells in a lineage over each time interval. In WT, cell division is localized more densely at the distal half of the sepal at 0-24 hrs, then and progresses proximally towards the base over time (Fig 4A, Fig S2G-H). This common spatial pattern is called a basipetal gradient (Hervieux et al, 201). Interestingly, *lgo-2* (Figure 4B, S2I-J) and *LGOoe* (Figure 4C, S2K-L) cell division is also localized in a basipetal gradient, but with increased and decreased divisions respectively. Therefore, cell division rate does not affect the localization of cell division.

**Figure 4:**
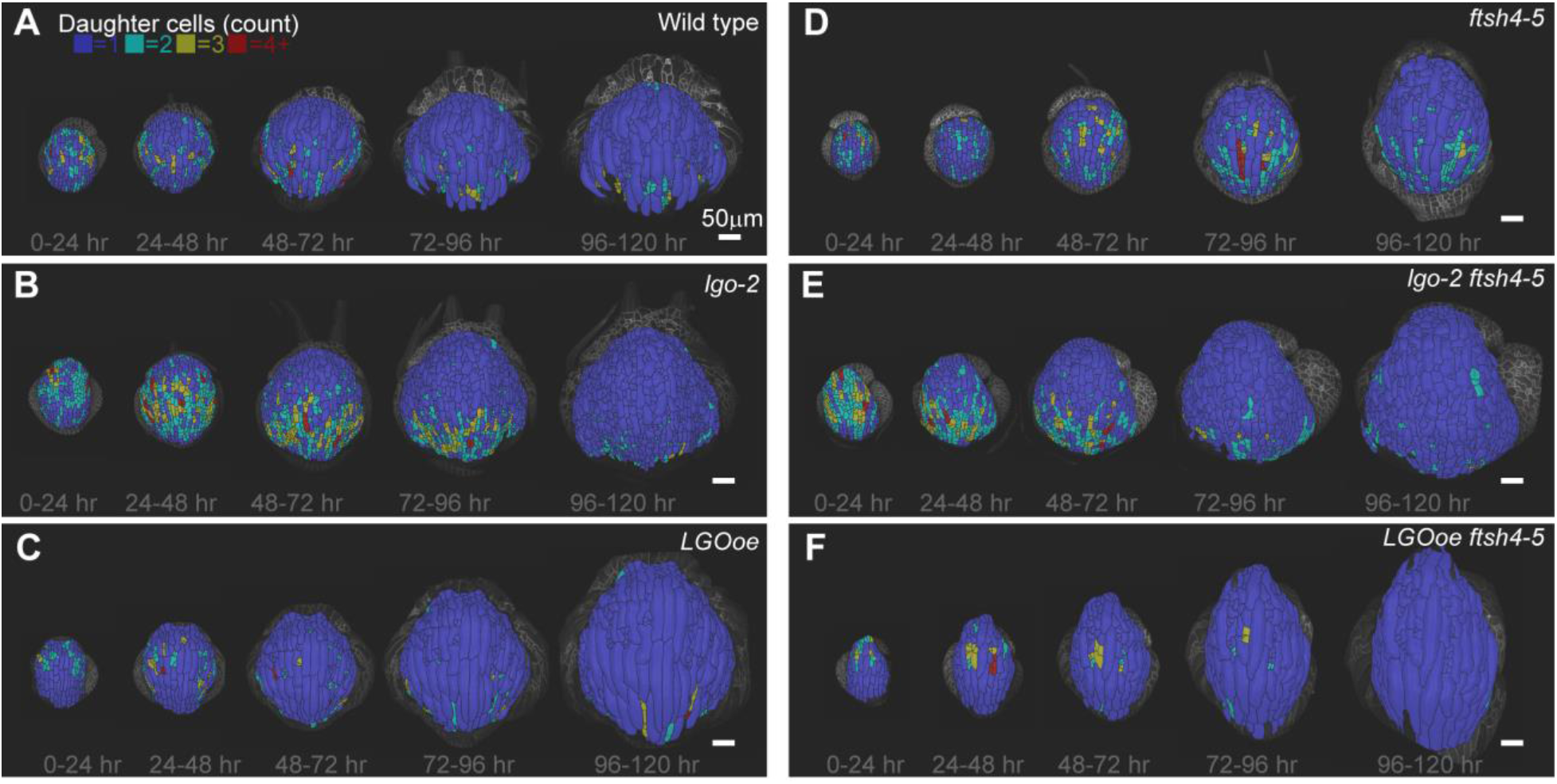
Cell division follows a basipetal gradient in WT, *lgo-2* and *LGOoe,* but not in *ftsh4-5*, *lgo-2 ftsh4-5*, and *LGOoe ftsh4-5*. (A-F) Heat maps of number of daughter cells per cell lineage over 24-hour intervals for WT (A), *lgo-2* (B), *LGOoe* (C), *ftsh4-5* (D), *lgo-2 ftsh4-5* (E), and *LGOoe ftsh4-5* (F). The lowest heat map value represents 1 cell per lineage, which means no division. The greatest heat map value represents 4 or more cells per lineage. The scale bar is 50µm. Heat maps are projected onto the later time point. Representative images from n=3 biological replicates. Additional replicates are available in Supplemental Figure S2.

In *ftsh4-5*, cell division has less tight distal localization at 0-24 hrs compared to WT. In later time points, cell division is more proximal, but localized in patches rather than a band that spans the sepal (Fig 4D. Fig S2M-N). Similar to *ftsh4-*5, division in *lgo-2 ftsh4-5* sepals does not show clear distal localization at the early time points. Neither *lgo-2 ftsh4-5* nor *LGOoe ftsh4-5* exhibit a band-like localization of cell division (Fig 3E,F, FigS2O-R). Our results indicate that changing the rate of cell division does not change the localization of cell divisions during development. However, *ftsh4-5* slightly alters the localization of cell division.

### Changing division rate does not change the spatial pattern of cell growth

To understand how dramatic differences in cell division rate are compensated to have little effect on sepal shape, we examined cell growth. The epidermal cell layer drives morphogenesis (Savaldi-Goldstein, 2007), so we focus our analysis on measuring cell growth in the epidermis. In WT, the highest rates of cell growth are localized distally at the 0-24 hr time interval, and then cell growth forms a band-like localization that becomes more proximal over each successive time interval (Fig 5A, S5A-B). This basipetal pattern matches previously described sepal growth (Hervieux et al, 2016, Hong et al, 2016). Localization of cell growth in *lgo-2* (Fig 5B, S5C-D) and *LGOoe* (Fig 5C, S5E-F) is remarkably similar to that of WT. Thus, cell division rate has little effect on sepal shape because localization of cell growth is unchanged.

**Figure 5:**
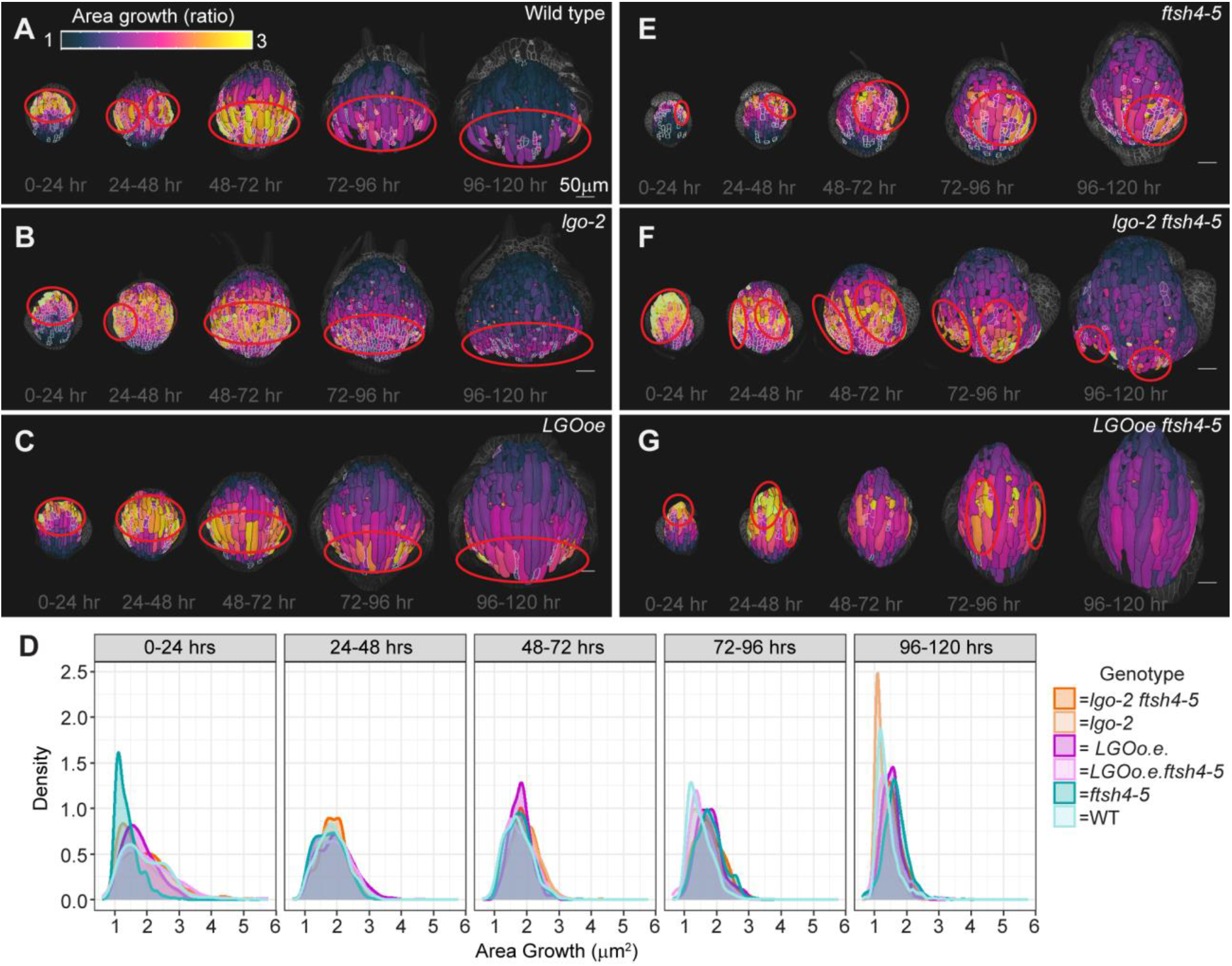
Cell growth follows a basipetal gradient which is preserved when cell division changes, but altered in *ftsh4-5, lgo-2 ftsh4-5*, and *LGOoe ftsh4-5.* (A-F) Heat maps of cell area growth over each 24-hour interval that are projected onto the later time point for WT (A), *lgo-2* (B), *LGOoe* (C), *ftsh4-5* (D), *lgo-2 ftsh4-5* (E), and *LGOoe ftsh4-5* (F). The heat map represents the change in ratio of cell area (cell area of later time point divided by cell area of earlier time point) and the scale is 1 to 3. The scale bar is 50µm. Daughter cells that result from a division over a given time interval area outlined in white. Localization of fast growth is marked by red outlines, and is band-like in WT, *lgo-2*, and *LGOoe* and patchy in *ftsh4-5*, *lgo-2 ftsh4-5*, and *LGOoe ftsh4-5*. Representative images from n=3 biological replicates. Additional replicates are available in Supplemental Figure S5. (D) Distribution of cell area growth for each time interval. Statistical analysis (multidimensional scaling) available in Supplemental Figure S6. Distribution of cell area growth related to the number of cell divisions in the lineage is available in Supplemental Figure S7.

However, the localization of cell growth in *ftsh4-5* differs from WT, *lgo-*2, and *LGOoe*. The sepals of *ftsh4-5* have localization of cell growth that appears patchy rather than band-like (Fig 5E, S5G-H). The patches can appear as faster growth that is spatially localized only on one side of the sepal or fast growth that persists for most of the time lapse. The localization of growth in *lgo-2 ftsh4-5* (Fig 5F, Fig S5I-J) and *LGOoe ftsh4-5* (Fig 5G, Fig S5K-L) matches that of *ftsh4-*5 in that it is patchy rather than band-like. Strikingly, some *lgo-2 ftsh4-5* sepals have especially clear boundaries between fast and slow growing patches. Therefore, growth localization is patchy and variable in the *ftsh4-5* background and is not affected by cell division rate.

Notably, the frequency distributions of cell growth rates of *ftsh4-*5, *lgo-2 ftsh4-*5, and *LGOoe ftsh4-5* do not differ from each other or from WT, *lgo-*2, and *LGOoe* (Fig 5D). Multidimensional scaling was used to compare the differences in the distributions of cell growth rates between genotypes and time points. This analysis reveals that all genotypes loosely cluster by time interval (developmental stage) but not by genotype (Fig S6) Thus, the localization of cell growth is disrupted in genotypes with a loss of robustness of shape, but not the rate of growth. WT, *lgo-2*, and *LGOoe*, which have robust sepal development, have the same spatial patterns of cell growth, whereas in *ftsh4-5*, *lgo-2 ftsh4-5* and *LGOoe ftsh4-5*, each replicate has a different spatial pattern of growth.

### Cell division and cell area growth colocalize but occur independently

We examined the relationship between cell area growth and cell division because both exhibit a basipetal gradient in WT, *lgo-*2, and *LGOoe.* To compare the localization of cell area growth and cell division, cells that formed from divisions were outlined on the cell growth heat maps. This reveals that fast cell growth and cell division colocalize to the same regions of the sepal, although the fastest growing cells are not necessarily the ones that divide in WT (Fig 5A, S5A-B), *lgo-2* (Fig 5B, S5C-D), and *LGOoe* (Fig 5C, S5E-F). Interestingly, fast cell area growth and cell division also colocalize in *ftsh4-5* (Fig 5E S5G-H)*, lgo-2 ftsh4-5* (Fig 5F, S5I-J), and *LGOoe ftsh4-5* (Fig 5G, S5K-L) despite the patchy localization patterns. Thus, across all genotypes, cells that divide in a given 24 hr interval tend to have greater cell area growth compared to cells that do not divide (Fig S7). The independence of cell growth from cell division in individual cells explains the robustness of sepal size and shape despite changes in cell division (Robinson et al, 2018) (Figure 1).

### Growth rate can vary within a cell

It is impressive that the basipetal pattern of growth is preserved in *LGOoe* considering giant cells span large vertical sections of the sepal. However, if giant cells had different amounts of expansion in different regions of the cells, this may facilitate the preservation of the normal localization of cell area growth. Different rates of growth within the same cell have been previously observed in the leaf (Elsner et al, 2012). To test this hypothesis, giant cells were artificially subdivided into multiple “cells.” Then cell area heat maps were created with the artificial “cells” outlined in white. In both WT (Fig 6A, S8A-B) and *LGOoe* (Fig 6B, S8C-D), different regions of the giant cells often had different amounts of area growth. Our data supports the conclusion that *LGOoe* basipetal growth is achieved by different growth rates within cells.

**Figure 6:**
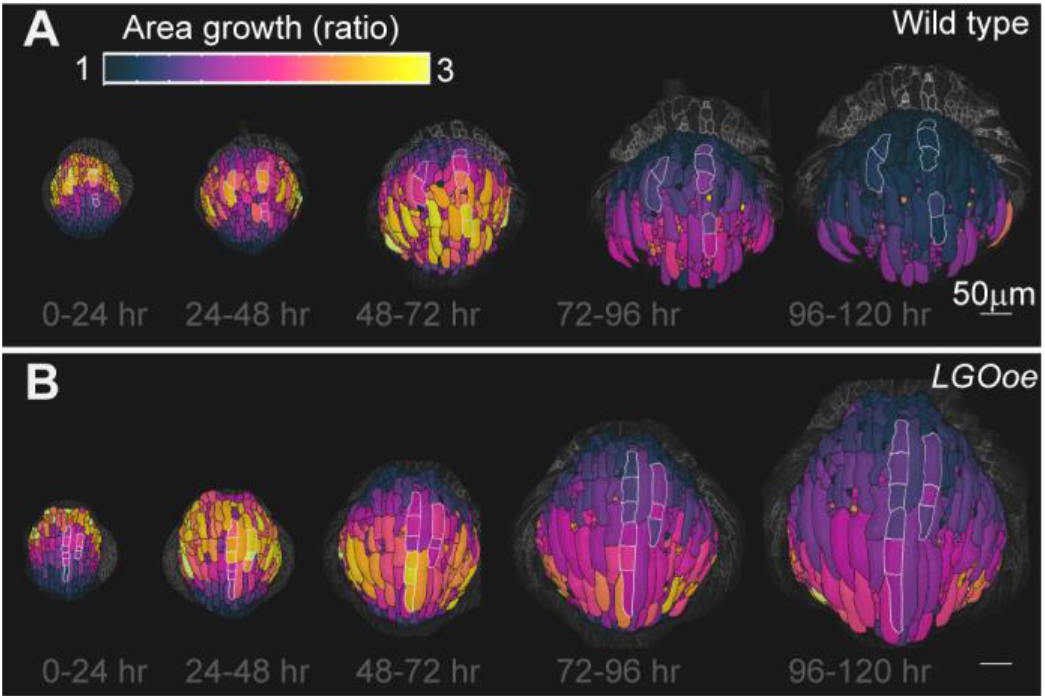
Cells can have regions with different growth rates in both WT and *LGOoe*. (A-B) Giant cells are artificially subdivided into multiple cells and outlined in white. The heat maps of cell area growth over each 24-hour interval are projected onto the later time point for WT (A) and *LGOoe* (B). The heat map represents the change in ratio of cell area (cell area of later time point divided by cell area of earlier time point) and the scale is 1 to 3. The scale bar is 50µm. Representative images from n=3 biological replicates. Additional replicates are available in Supplemental Figure S8.

### Spatiotemporal averaging of heterogeneous cell growth rate occurs when cell division is altered, but not in *ftsh4* mutants

We have previously proposed that spatiotemporal averaging of heterogeneous cell growth creates robustness in sepal shape (Hong et al, 2016). Previously we have assessed spatiotemporal averaging only in growth direction, not in growth rate. Here we assess spatiotemporal averaging in growth rate by measuring cumulative growth over three days (spanning the 24 hr to 96 hr time point). If the heterogeneous cell growth rates average, we expect the cumulative growth of cell lineages in the sepal should become more uniform. Since spatiotemporal averaging occurs superimposed on the basipetal growth gradient, we expect that this averaging will generate a band of relatively uniform fast cell growth in the middle of the sepal surrounded by bands of slower cell growth at the tip and base. That is what we observe in WT; cumulative three-day growth heat maps have a band of relatively uniform growth across the medial-lateral axis in all three replicates (Fig 7A). Cells that are at the same location along the proximal distal axis have similar amounts of growth. These bands of uniform cumulative growth are consistent with the model that underlying heterogeneity in growth rates average over time into even growth across the organ. Cumulative growth rates form uniform bands in lgo-2 (Fig 7B) and LGOoe (Fig 7C) similar to WT, suggesting that spatiotemporal averaging of cell growth occurs despite changes in the cell division rate.

**Figure 7:**
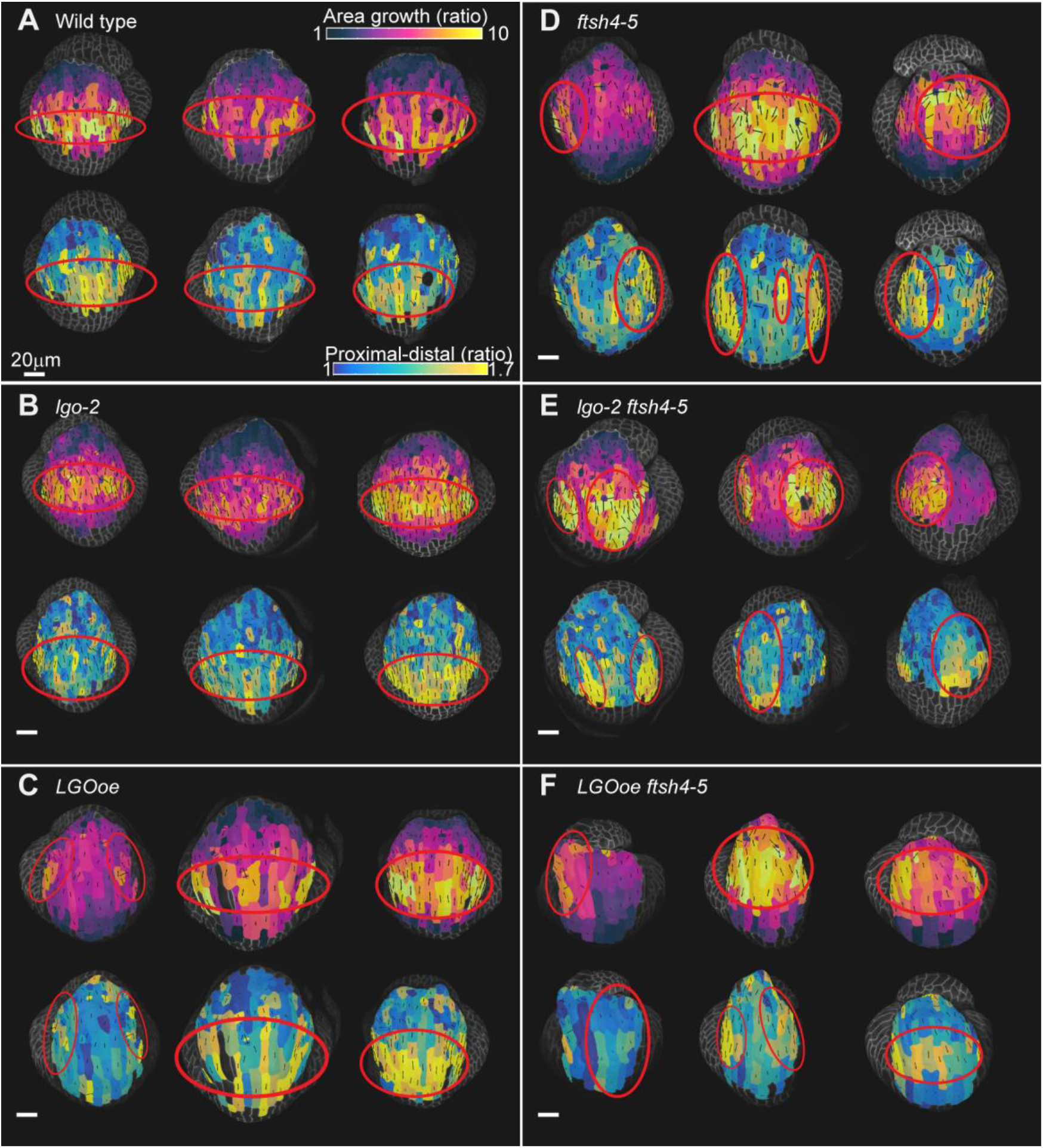
Cell growth rate and direction averages to be uniform across the organ in WT, *LGOoe*, and *lgo-2* but do not average in *ftsh4-5*, *lgo-2 ftsh4-5*, and *LGOoe ftsh4-5.* (A-F) Heat maps from 24-hour time point to 96-hour time point for cell area growth (top rows) and ratio of proximal-distal cell growth to medial-lateral cell growth (bottom rows) projected on the earlier time point for WT (A), *lgo-2* (B), *LGOoe* (C), *ftsh4-5* (D), *lgo-2 ftsh4-5* (E), and *LGOoe ftsh4-5* (F). Three replicates are shown for each genotype, and each replicate has heat maps of both measures. The scale bar is 50µm. The principal directions of cell growth are overlaid on the heat maps as black lines that are oriented in the direction that each cell had the most growth and have a length that corresponds to the magnitude of the ratio of growth parallel to the principal direction of growth to the growth perpendicular to the principal direction of growth. Top rows: The heat map represents the change in ratio of cell area (cell area of later time point divided by cell area of earlier time point) and the scale is 1 to 10. Bottom rows: A proximal distal axis was defined, and the heat map represents the ratio of cell growth parallel to the axis divided by perpendicular to the axis. The lowest heat map value is 1, which represents equal amounts of growth along both axes. The highest heap map value is 1.7, which represents 1.7x more proximal-distal growth than medial-lateral growth. Red circles mark the region of the sepal with greater area growth (top rows) and greater proportion of proximal-distal growth (bottom rows). Quantification of the relationship between growth rate and proximal distal growth orientation is available in Supplemental Figure S9.

In contrast, cumulative cell area growth over 3 days in ftsh4-5 sepals usually does not appear uniforms; instead, the patchy growth accumulates into more pronounced patches of fast and slow growth (Fig 7D), suggesting a lack of spatiotemporal averaging of cell growth over time. Although one ftsh4-5 replicate appears less patchy, it has a large region in the center that grows even faster than WT, suggesting that the center of the sepal has excess growth. Every ftsh4-5 sepal has a different cumulative growth pattern, which fits with the variability of sepal size and shape phenotype. Cumulative cell area growth is similarly patchy in lgo-2 ftsh4-5 (Fig 7E) and LGOoe ftsh4-5 (Fig 7F). In many of the replicates, a patch of slower growing cells persists over time and intervenes between patches of faster growing cells. Examining the same sample over consecutive time points (Figure 5E,F, G) and cumulatively (Figure 7D,E F), shows that slower growing cells in ftsh4-5, ftsh4-5 lgo-2, and ftsh4-5 LGOoe mutants persist in their slow growth over consecutive time points, such that these same cells appear as the cumulative slow growth patch. In comparison, a cell that is growing slowly in wild type, lgo-2, or LGOoe is often growing faster in the next time point, allowing the growth rate to average over time, producing a moderate overall cumulative cell growth rate. Together, our data suggest that in ftsh4, cell growth patterns are stabilized in time, resulting in the patchy cumulative growth indicating a loss of averaging. This cumulative patchy growth is expected to cause unpredictable, uneven expansion of the organ. Furthermore, changing the rate of cell division does not alter the loss of spatiotemporal averaging in ftsh4-5.

### Cell growth direction is organized in even growth and disorganized in patchy growth

Cell area growth rate measures the magnitude of growth. However, the direction of cell expansion is also an important consideration for development of shape. Previously, it has been shown that direction of cell growth averages over time to be aligned with the proximal-distal axis in WT sepals, and not in *ftsh4-5* sepals (Hong et al, 2016). To test whether cell division rate affects averaging of growth direction, we examined the cumulative principal direction of growth over 3 days, which is the primary direction that the cell or cell lineage expands over the time interval. In WT, *lgo-2* and *LGOoe*, the cells in the band of fast, even growth have principal directions of growth that appear aligned to each other and to the proximal-distal (base to tip) axis of the sepal (Figure 7A-C). A trichome is present in one WT replicate, which is known to cause neighboring cells to alter growth (Hervieux et al, 2017), and explains the unaligned growth directions in those cells. However, in *ftsh4-5, lgo-2 ftsh4-5,* and *LGOoe ftsh4-5*, the patches of fast growth have cells with variable directions instead of directions aligned with the proximal distal axis (Figure 7D-F). Therefore, genotypes with robust shape have even tissue expansion in an organized direction whereas genotypes with loss of robustness of shape have patchy tissue expansion in variable directions. Cell division rate does not affect the averaging of cell growth direction.

### Even growth, but not patchy growth, overlaps with proximal-distal elongation

Although WT, *lgo-2,* and *LGOoe* have decreased variability in cell growth direction in regions of faster growth, *ftsh4-5*, *lgo-2 ftsh4-5*, and *LGOoe ftsh4-5*, have increased variability in growth directions between cells in faster growing patches. To better compare the spatial localizations of proximal-distal growth with the cell growth rates, 3-day principal directions of growth were converted to heat map values. A value of 1 would signify equal amounts of proximal-distal and medial-lateral cell growth (isotropic growth). A heat map value greater than one indicates that proximal-distal growth is greater than medial-lateral growth. In WT, *lgo-2*, and *LGOoe*, the greatest amount of proximal-distal growth is localized in a band across the center of the sepal (Figure 7A-C) and colocalizes with fast cell growth. Further, greater cell growth is correlated with greater proximal-distal elongation (Fig7SA-C). This trend is different in *ftsh4-5*, *lgo-2 ftsh4-5*, and *LGOoe ftsh4-5*. Instead, the cells with the greatest proximal-distal growth do not colocalize with fast cell growth, and instead colocalize with patches of slow growth (Figure 7D-F). Further, *ftsh4-*5, *lgo-2 ftsh4-*5, and *LGOoe ftsh4-5* have a weaker correlation between growth rate and proximal-distal growth (Figure S9D-F). Therefore, *ftsh4-5* causes a loss of colocalization of fast growth and proximal distal elongation. The coordination of fast growth and proximal-distal growth in WT, *lgo* and *LGOoe* is likely responsible for the uniform sepal size and shape. On the other hand, variability in temporal and spatial location of growth and variable growth directions should both lead to variable organ size and shape.

## Discussion

Sepal development results in uniform organ size and shape in WT, indicating that development is robust. The development of organs has been shown in many situations to be robust to changes in cell division, a phenomenon known as compensation. To understand how compensation occurs and makes sepal morphogenesis robust to changes in cell division, we time lapse imaged a mutant with increased cell division rate (*lgo-2*), a transgenic plant with decreased cell division (*LGOoe*), a mutant with variable organ size and shape (*ftsh4-5*), and double mutants. We confirm that sepal development is robust to changes in cell division. Further, the loss of robustness in *ftsh4-5* is not affected by changing cell division rate with *lgo-2* or *LGOoe*. We find that cell growth localization and growth direction are not affected by the cell division rate, which explains robustness to changes in cell division. The change in cell division rate without changing growth automatically generates the change in cell size observed during compensation. For instance, increasing the division rate while maintaining the growth rate automatically generates more smaller cells. Thus, our imaging reveals one mechanism through which compensation occurs.

It has been previously proposed that WT sepal development is robust to heterogeneity in cell growth rate through spatiotemporal averaging. Here we find further evidence of spatiotemporal averaging, because cell growth rates form uniform bands of growth across the sepal. We find that the cells in the bands with the highest growth rates also align their growth to the proximal distal axis. Thus, each of these WT developing sepals are expanding evenly and in the same direction, which should cause them to have similar final sepal shapes. We find the same evidence of spatiotemporal averaging in *lgo-2* and *LGOoe* as in WT, revealing that sepal development is robust to changes in cell division both because cell division does not change the growth patterns and because those growth patterns still undergo spatiotemporal averaging.

Spatiotemporal averaging of growth rate and direction is disrupted in the *ftsh4-5, lgo-2 ftsh4-5*, and *LGOoe ftsh4-5* mutants, which also have a loss of robustness of shape. Instead, the tissue accumulates patches of fast and slow growth, and the fastest growing cells are often not elongating in the proximal-distal direction. The patchy and disorganized growth direction suggests that these sepals will not be uniform in shape. These results are consistent with previous modeling in which high spatial correlation in growth rate causes regions of the tissue to grow different amounts, resulting in variable organ size and shape (Hong et al, 2016). Therefore, averaging heterogeneity during development is necessary for robust development of shape.

### Robustness of shape is preserved despite changes in cell division

WT, *lgo-2*, and *LGOoe* have similar variability in mature sepal shape, indicating that robustness of sepal shape is not influenced by cell division rate. Further, the loss of robustness in *ftsh4-5* is not affected by changing cell division rate with *lgo-2* or *LGOoe*. The ability to preserve robust development of shape despite cell division stems from the preservation of the spatial localization of growth despite changes in cell division rate. Growth occurs in a basipetal gradient in WT, *lgo-2*, and *LGOoe*. Further, growth rates can vary at the subcellular level, which may assist *LGOoe* in replicating the basipetal growth gradient with mostly large cells that span different growth rate regions.

Robustness is preserved in *lgo-2* and *LGOoe*, which are cell division mutants which exhibit compensation between cell size and cell number during the proliferative stages of organ development. However, some mutants with decreased cell number exhibit compensation after the proliferative phase of organ growth is over (Ferjani et al, 2007). This suggests that additional mechanisms to preserve robustness can occur after the proliferative phase of organ development in addition to the spatiotemporal averaging mechanism we have found that occurs during the proliferative phase.

### Reproducible cell growth localization is associated with uniform sepal size and shape

A characteristic of the *ftsh4-5* phenotype is variability in both final sepal size and shape and the localization of growth during development. The variability in the *ftsh4-5* phenotype allows for evaluation of robustness of shape rather than development of a particular shape. This means that different replicates of *ftsh4-5* were variable compared to each other whereas wild type replicates were more consistent. Despite variability in sepal size and shape, a lack of averaging of heterogeneity, patchy growth rate, and disorganized growth direction, were characteristic of all *ftsh4-5* sepals that were imaged. Therefore, these characteristics are indicative of a loss of robust development rather than development of a certain size or shape.

### Spatiotemporal averaging of heterogeneity occurs in vivo, and is preserved despite changes in cell division rate

Previously it was found that WT has more spatiotemporal heterogeneity than *ftsh4-5* in cell area growth rates (Hong et al, 2016). Modeling predicted that if a 2D growing shape has regions with different specified growth rates, and if the growth rates frequently change, then the heterogeneity present at single time points will average over time and cause all parts of the shape to have equal growth. There is less spatial and temporal heterogeneity in *ftsh4-5* cells growth during sepal development (Hong et al, 2016). Accordingly, the model was also modified to decrease spatial and temporal heterogeneity in growth by removing changes in growth rate and increasing the size of regions with a specified growth rate. This causes some parts of the shape to grow more than others, and there is variability in the final shapes when the model is run multiple times (Hong et al, 2016). Here, we time lapse image sepal development for a longer time period, which allows us to understand the long-term implications of heterogeneity and spatiotemporal averaging *in vivo.* The patchy localization of growth observed in *ftsh4-5*, *lgo-2 ftsh4-5*, and *LGOoe ftsh4-5* is remarkably similar to the model. Therefore, the time-lapse imaging here supports the findings of the model that spatiotemporal averaging of heterogeneous growth leads to robust sepal development. Further, spatiotemporal averaging is preserved when cell division rate changes.

### Colocalization of spatiotemporal averaging of growth rate and growth direction

Previously, it was found that heterogeneity in growth direction averaged over time in WT and not in *ftsh4-5* (Hong et al, 2016). Here we find that averaging of growth direction occurs in the sepal cells with the highest growth rates, which results in fast growth in an organized (proximal-distal) direction. This coordination between growth rate and direction will lead to the development of a uniform, reproducible shape. There is a different relationship between growth rate and direction in the *ftsh4-5* background. Instead, the fastest growing cells have growth directions that are not growing in the proximal-distal direction and are not aligned within the organ. It remains to be determined whether disorganized growth direction is independent of patchy growth, or if patchy growth rotates these cells relative to the rest of the organ resulting in disorganized growth direction. Persistent patches of fast growth with variable growth direction lead to variable organ shape.

## Methods

### Plant material

Accession *Col-0* plants are used as wild-type and all mutants are in *Col-0* background as well. Isolation of the *ftsh4-5* mutant is described in Hong et al, 2016. *ATML1p::LGO* (*LGOoe*) is from Roeder et al 2010. The membrane marker *p35S::mCitrine-RCI2A* was crossed into *lgo-2*, *LGOoe* and *lgo-2 ftsh4-5*. *35S::mCitrine-RCI2A* was transformed into *LGOoe ftsh4-5* due to silencing. The epidermal specific membrane marker *ML1::mCitrine-RCI2A* was used in *ftsh4-5* plants due to silencing.

### Genotyping

The *lgo-2* mutation can be PCR genotyped with the primers CTTCCCTCTCACTTCTCCAA, CCGAACACCAACAGATAATT, and TTGGGTGATGGTTCACGTAGTGGG. The WT band is 546 base pairs and the *lgo-2* band is 753 base pairs. The *ftsh4-5* mutation can be PCR genotyped with the primers AGAAAGGACTCACTTTAAAGAACAGCCATG and TCCTCTGTCCTCGATAAGAGCTCC followed by digesting the product with Nco1 which produces a WT band of 103 base pairs and a *ftsh4-5* band of 124 base pairs. The *LGOoe* plants are easily distinguished by their phenotype of curled leaves.

### Images of phenotypes and sepal shape variability quantification

Photographs of mature flowers and flower buds were taken with either a Canon A610 or an Excelis 4K camera mounted on a Zeiss Stemi stereomicroscope. Sepals of mature flowers were dissected, placed on a black background, flattened under a slide, and photographed using the Canon A610 camera mounted to a dissecting microscope. Python programs, as described and available in Hong et al, 2016, were used to trace the outline of the sepal shapes and converted into contours of the shape and measurement of area. Shapes were normalized by size and then the variability of shape was compared between genotypes.

### Microscopy and Image Analysis

Inflorescences were dissected and mounted in apex culture media Hamant et al, 2014. Media contained 2.3 g/L Murashige and Skoog, 1% sucrose, 0.1% MES pH=5.8 media supplemented with vitamins (final concentration of 100 µg/ml myoinositol, 1 ng/ml nicotinic acid, 1 ng/ml pyridoxine hydrochloride, 1 ng/ml thiamine hydrochloride, 2 ng/ml glycine), plant preservative mixture from Plant Cell Technology which was used as 1000X stock, and 1.2% agarose. Plants then grew in 16 hr light 8hr dark conditions on the media and were imaged with a Zeiss LSM710 confocal microscope once every 24 hrs for 6 days. A 20X water dipping objective with an NA of 1.0 (W Plan-APOCHROMAT 20x.1.0 DIC (US) VIS-IR) was used. A 514 laser with a power of 5% was used for excitation. The voxel size was x=0.4151, y=0.4151, z = either 1.5 μm or 0.5 μm. The wavelengths detected were 519-622 nm. The zoom was 1. The pinhole was 2.17 airy units=3.4 μm.

MorphoGraphX was used for image processing (Barbier de Reuille et al., 2015; Strauss et al., 2022). One of two methods was used to detect the surface. Method one begins by trimming voxels of a trichome if there is one was present, then Stack/Filters/Gaussian Blur Stack(X sigma=1, Y sigma=1, Z sigma=1), Stack/Morphology/Edge Detect (Threshold=3000-10000, Multiplier=2.0, Adapt= 0.3, Fill Value= 3000) to find the surface, then either Stack/Morphology/Closing (X Radius=1-10, Y Radius= 1-10, Z Radius=1-10) or Stack/Morphology/Fill Holes (X Radius= 1-10, Y Radius=1-10, Threshold=10000, Depth= 0, Fill Value= 30000) to fix any holes and then manually trimming voxels of adjacent organs or empty space that was filled by the Fill Holes function. Method two made the surface using Stack/Lyon/Init Level Set (Up Threshold=2-10, Down Threshold=2-10), Stack/Lyon/Level Set Evolve (Default settings except View=5 and cancel after 5 to 15 rounds), Stack/Morphology/Edge Detect Angle, and Stack/Morphology/Closing (X Radius=1-10, Y Radius= 1-10, Z Radius=1-10) and manually trimming voxels of adjacent organs. Then the mesh was created with the processes Mesh/Creation/Marching Cubes Surface (Cube size= 5.0, Threshold=20000), then 2-4 rounds (smaller meshes had 3-4 and larger meshes had 2-3) of Mesh/Structure/Subdivide and Mesh/Structure/Smooth Mesh (Passes= 10, Walls Only= No). Then the mesh was segmented by projecting a 2 μm depth interval of the signal using Mesh/Signal/Project Signal (Min Distance= 2-8, Max Distance= 4-10) followed by Mesh/Segmentation/Watershed Segmentation (Steps= 50000). Lineage tracking was done by loading meshes for consecutive time points into mesh 1 and mesh 2 spots, overlapping mesh 1 (check scale box and increased size) and mesh 2 using the shapes of the cell lineages, then either manually or semi-automatically assigning parent labels. Parent labels were checked for errors by running Mesh/Cell Axis/PDG/Check Correspondence on the earlier time point. To make sure there were no cells on the periphery that were parent tracked, but were partially cut off by the edge of the images in the later time point, Mesh/Heat Map/Heat Map Classic (change map checked, decreasing) was run, and Mesh/Heat Map/Heat Map Select (Lower=0, Upper=.999) was used to highlight cells that had “shrunk.” If the cells were at the edge of the segmentation, they were assumed to be a segmentation error, and deleted. Then Mesh/Lineage Tracking/Save Parents was run on the later time point to save the parent labels as a csv file. Then Mesh/Lineage Tracking/Load Parents was run to load the csv file that was just created and then meshes were saved with the parent labels.

To make the 120-hour cumulative cell division heat maps, csv files specifying parent labels for the 0 hr to 120 hr time points were created from the 24 hr parent label csv files using a python script to do multi-step lineage tracking as described in Hong et al, 2016. Then the corresponding parent labels were loaded onto the later time point using Mesh/Lineage Tracking/Load Parents. Mesh/Lineage Tracking/Heat Map Proliferation and Mesh/Heat Map/Heat Map Set Range (Min=1, Max=15) were run on the later time point to make the heat map and they were saved as csv files using Mesh/Heat Map/Heat Map Save.

To make the cell area heat maps Mesh/Heat Map/Geometry/Area and Mesh/Heat Map/Heat Map Set Range (Min=0 Max=4000) were run on each mesh and csv files were saved with Mesh/Heat Map/Heat Map Save.

To make the 24 hr proliferation heat maps Mesh/Lineage Tracking. Mesh/Lineage Tracking/Heat Map Proliferation and Mesh/Heat Map/Heat Map Set Range (Min=1 Max=4) were run on the later time point of each 24 hr interval. Mesh/Heat Map/Heat Map Save was run to save the csv files.

To make the cell area heat maps with outlined divisions, Mesh/Heat Map/Heat Map Classic (changed map checked, decreasing) was used to create the cell growth heat map, and then it was saved as a csv file using Mesh/Heat Map/Heat Map Save. Then the 24 hr proliferation heat maps were loaded onto the later time point of the 24 hr interval using Mesh/Heat Map/Heat Map Load. The Mesh/Heat Map/Heat Map Select (Lower Threshold=2, Upper Threshold=7) was used to outline the cells that had divided at least once. Then Mesh/Heat Map/Heat Map Load was used to load the cell growth heat map, and Mesh/Heat Map/Heat Map Set Range (Min=1, Max=3) was used to set the scale.

To artificially subdivide the giant cells, a few giant cells were chosen to be deleted from the mesh based on nearby junctions that would be helpful landmarks. The cells were manually seeded as multiple cells using nearby junctions as landmarks, parent tracked as described above. Cell labels were outlined using Mesh/Selection/Select Labels (add the labels of the cells). Cell growth heat maps were created using Mesh/Heat Map/Heat Map Classic (changed map checked, decreasing, use manual range 1-3) on the later time point.

To make the 3-day area growth and principal directions of growth, csv files specifying parent labels for the 24 hr to 96 hr time points were also made using a python script for multi-step lineage tracking as described in Hong et al, 2016. Then the corresponding parent labels were loaded onto the later time point using Mesh/Lineage Tracking/Load Parents. Mesh/Heat Map/Heat Map Classic (changed map checked, decreasing, use manual range 1-10) was run on the later time point, and Mesh/Heat Map/Heat Map Save was used to save the heat map as a csv file. Then Mesh/Cell Axis/PDG/Check Correspondence and was saved as a csv file using Mesh/CellAxis/PDG/Compute Growth then Mesh/Cell Axis/Cell Axis Save. Then the process /Unselect was run on mesh 1 or the mesh was reloaded, and the csv file was reloaded using Mesh/Cell Axis/Cell Axis Load. Then Mesh/Cell Axis/PDG/Display Growth Directions (Show Axis=StrainMax, Color+=black, Line Width=5, Line Scale=2) was used to display the principal directions of growth, Mesh/Heat Map/Heat Map Load was used to load the cell growth heat map that was saved as a csv file and Mesh/Heat Map/Heat Map Set Range (Min=1, Max=10) was used to adjust the scale.

To make the heat maps of proximal-distal growth, a custom axis was created from a heat map of distance from cells at the tip of the sepal, then the proportion that the principal directions of growth were aligned with the custom axis was used to create heat map values. First, to create the distance heat map, cells were manually selected and then the process Mesh/Heat Map/Measures/Location/Cell Distance was run. This was often repeated with different cells selected until the heat map appeared to measure distance accurately instead of creating a gradient that was curved or crooked. Then the heat map was saved as a csv file with Mesh/Heat Map/Heat Map Save. Then the principal directions of growth were loaded with Mesh/Cell Axis/Cell Axis Load. Then the distance heat map was loaded with Mesh/Heat Map/Heat Map Load. Then Mesh/Cell Axis/Custom/Create Heatmap Directions (Project Directions…=Yes, Normalize=no) and Mesh/Cell Axis/Custom/Smooth Custom Directions (Weight by Cell Area=Yes, Project Directions…=Yes) were used to create the axis from the distance heat map. Then Mesh/Cell Axis/PDG/Display Growth Directions (Heatmap=Aniso Custom, ScaleHeat=Manual, Heat min=1, Heat max=1.7, Show axis=StrainMax, color+=black, Line Width=5, Line Scale=2) was used to create a heat map of the ratio of the amount that the principal directions of growth were parallel with the custom axis to the amount that they were perpendicular to the custom axis.

### Analysis and Statistics

One-way ANOVA with genotype as a variable and Tukey tests were used to test for differences in shape variability, cumulative proliferation, and number of nondividing cells. Kendall rank correlation was used to test if there was a relationship between area growth and proximal-distal growth in each genotype. Wasserstein tests were used to create the principal coordinate analysis plots. R scripts are available at DOI: 10.17605/OSF.IO/7NMK. The R version 4.2.2 was used for analysis. The R packages used were ggplot2_3.4.1, tidyr_1.3.0, stringr_1.5.0, dplyr_1.1.0, twosamples_2.0.0, RColorBrewer_1.1-3, ggsci_2.9, ggthemes_4.2.4, ggplot2_3.4.1.

## Acknowledgements

We thank Lilan Hong, Shuyao Kong, Avilash Singh Yadav, Maura Zimmermann, and Michelle Heeney for helpful discussions and comments on the manuscript. We thank Cornell Statistical Consulting Unit (Matt Thomas) for helping with analysis of cell area, proliferation, and cell growth in R. We thank Mingyuan Zhu and Richard Smith for expert image analysis advice. We thank Olivier Hamant, Arezki Boudaoud, Corentin Mollier, Ya Min for helpful discussions. We thank Avilash Singh Yadav for technical help with an experiment.

## Competing interests

The authors declare they have no competing interests.

## Funding

Research reported in this publication was supported by the National Institute of General Medical Sciences of the National Institutes of Health under Award Number R01GM134037. The content is solely the responsibility of the authors and does not necessarily represent the official views of the National Institutes of Health.

## Data Availability

Data for this project is available at DOI: 10.17605/OSF.IO/7NMK3

## Author Contributions

Conceptualization: AHKR and IB

Genetics to create plant lines: IB and FKC

Experiments and image processing: IB

Shape variability analysis and multidimensional scaling: C-BL

Data Analysis: IB

Writing: IB

Revising and editing: IB. AHKR, C-BL, FKC

## Notes

### Competing Interest Statement

The authors have declared no competing interest.

### Summary of Updates

Supplemental figures are added because they did not automatically upload.

