## Supplemental Figures for "Robust organ size in Arabidopsis is primarily governed by cell growth rather than cell division patterns"

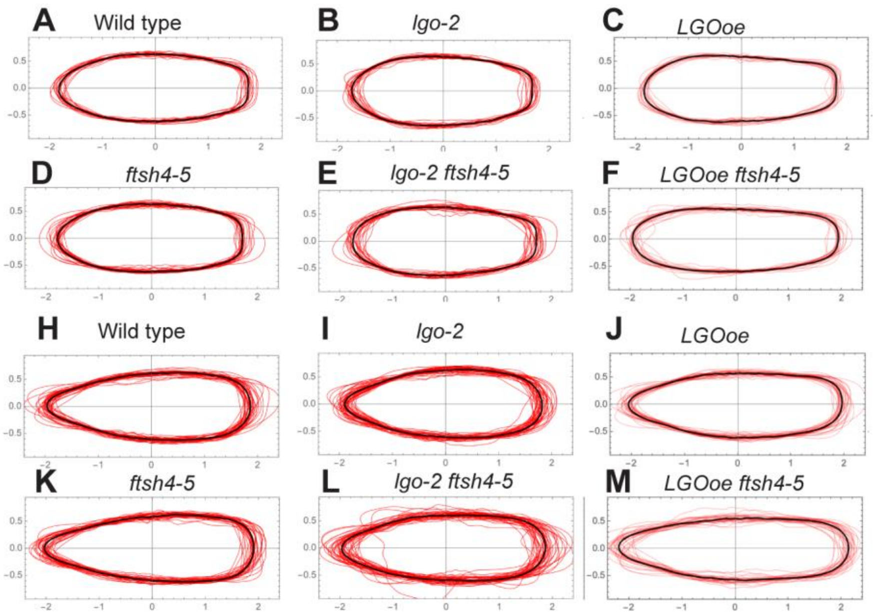


Figure S1: **Organ shape is robust to changes in *LGO* expression level, but not in *ftsh4-5*.** (A-F) Contours of mature inner (adaxial) sepals normalized by size (red lines) and the average sepal shape (black line). n=20 (WT), 22 (*lgo-2*), 22 (*LGOoe*), 26 (*ftsh4-5*), 25 (*lgo-2 ftsh4-5*), 22 (*LGOoe ftsh4*) (H-M) Contours of mature lateral sepals normalized by size (red lines) and the average sepal shape (black line). n=39 (WT), 42 (*lgo-2*), 34 (*LGOoe*), 35 (*ftsh4-5*), 44 (*lgo-2 ftsh4-5*), 30 (*LGOoe ftsh4*)

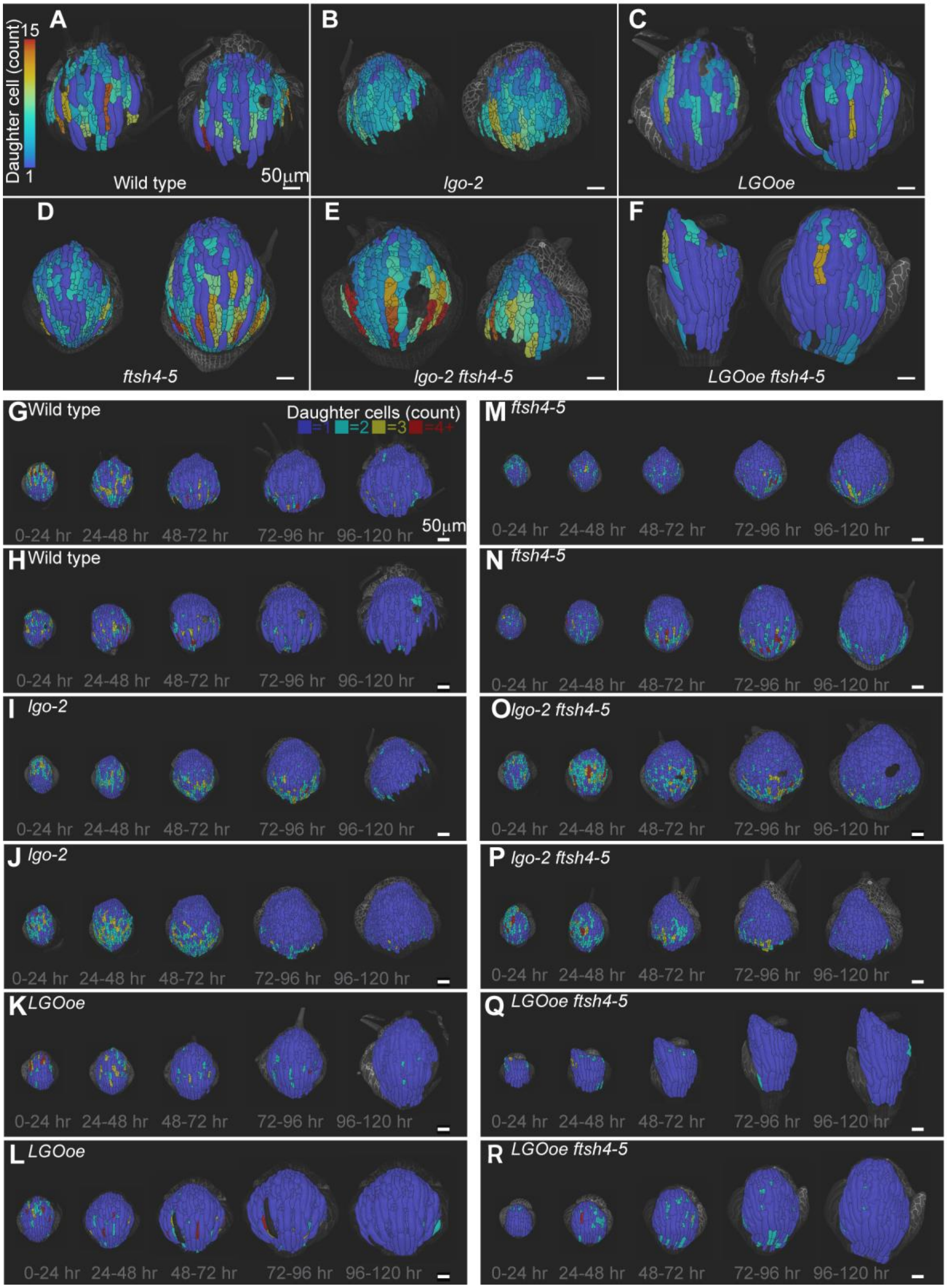


Figure S2: **Cell division rate is decreased by *LGO overexpression* (*LGOoe*) and *LGOoe ftsh4* and is increased in *lgo-2* and *lgo-2 ftsh4-5*.** (A-F) Heat maps of number of daughter cells per lineage using lineage tracking from 0-hour time point to 120-hour time point that are projected onto the 120-hour time point for remaining two replicates each of WT (A), *lgo-2* (B), *LGOoe* (C), *ftsh4-5* (D), *lgo-2 ftsh4-4* (E), and *LGOoe* (F). The heat map scale is 1 to 15 daughter cells and the scale bar is 50µm. (G-R) Heat maps of number of daughter cells per lineage over 24-hour intervals for remaining two replicates of WT (G-H), *lgo-2* (I-J), *LGOoe* (K-L), *ftsh4-5* (M-N), *lgo-2 ftsh4-4* (O-P), and *LGOoe* (Q-R). The lowest heat map value represents 1 cell per lineage, which means no division. The greatest heat map value represents 4 or more cells per lineage. The scale bar is 50µm. Heat maps are projected onto the later time point. Related to Figures 2 and 4.

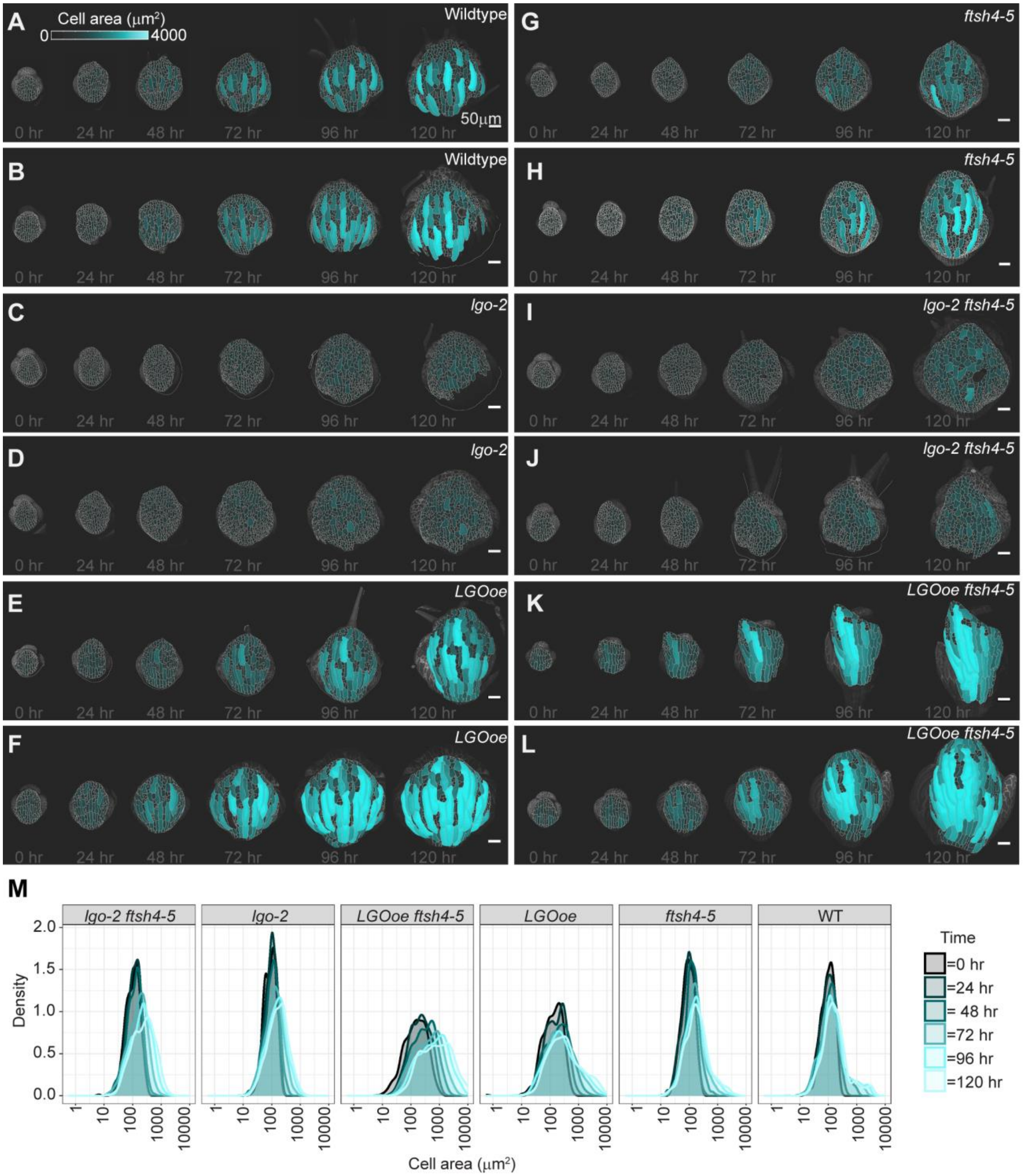


Figure S3**: Cell sizes remain smaller in *lgo-2,* and *lgo-2 ftsh4-5,* and become progressively larger in *LGOoe* and *LGOoe ftsh4*.** (A-L) Heat maps of cell area at each image time point for remaining two replicates of WT (A-B), *lgo-2* (C-D), *LGOoe* (E-F), *ftsh4-5* (G-H), *lgo-2 ftsh4-4* (I-J), and *LGOoe* (K-L). The heat map scale is 0 to 4200µm$^2$ and the scale bar is 50µm. (M) Density plots showing how the distribution of cell areas for each genotype changes over time. Related to Figure 3 and Supplemental Figure S4.

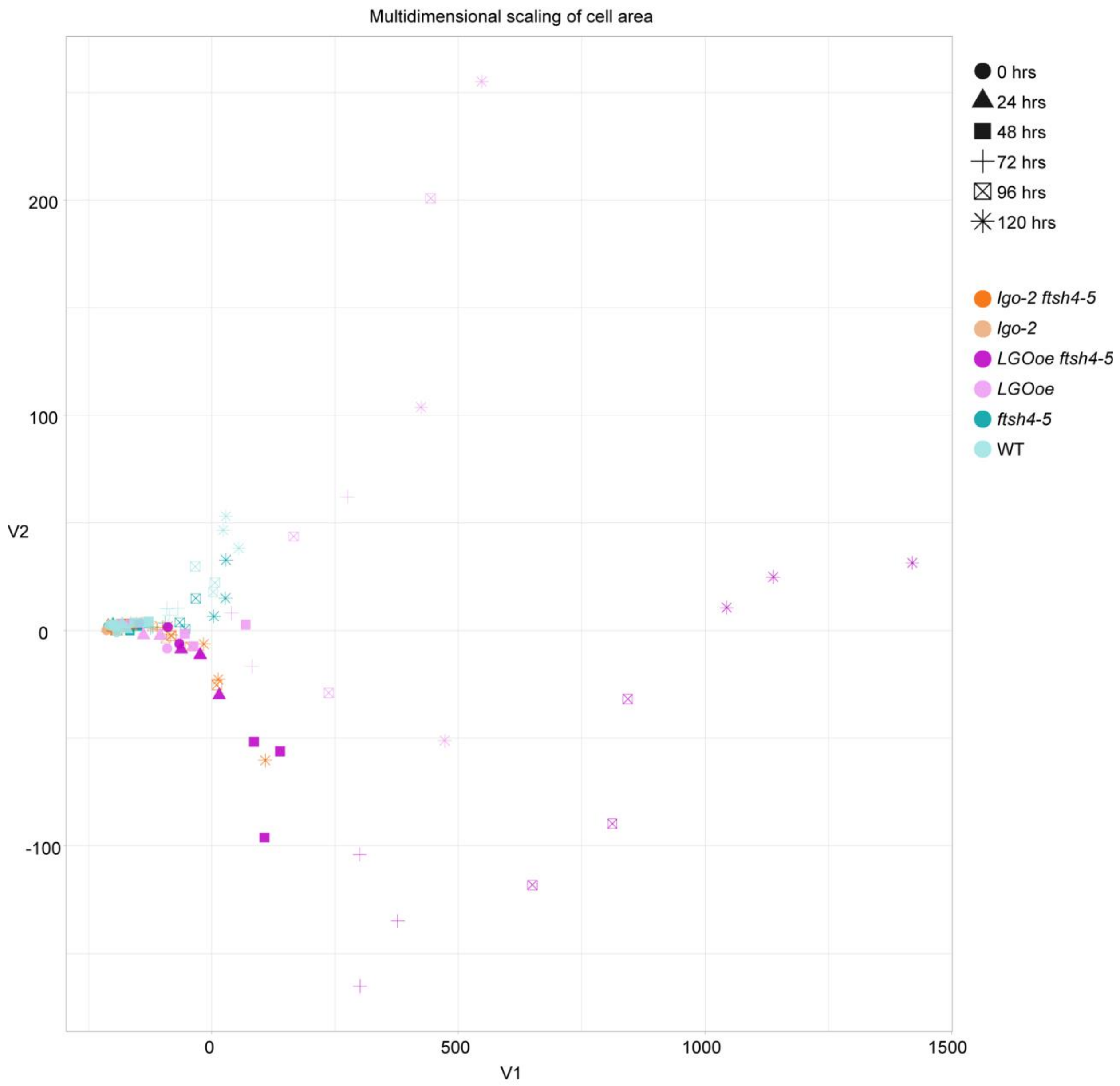


Figure S4: **Cell size diverges over time due to differences in cell division.** Multidimensional scaling (also called principal coordinate analysis) is used to represent the distributions of cell areas for each genotype and time interval in two-dimensional space. Increased distance between points indicates larger differences in the cell size distributions whereas points clustered together indicates that these cell size distributions are similar. Colors and shapes represent genotypes and time intervals, respectively. Note the cell size distributions of all genotypes cluster at the 0 hr and 24 hr time points, but then spread out based on the cell division rate. Related to Figure 3 and Supplemental Figure S2.

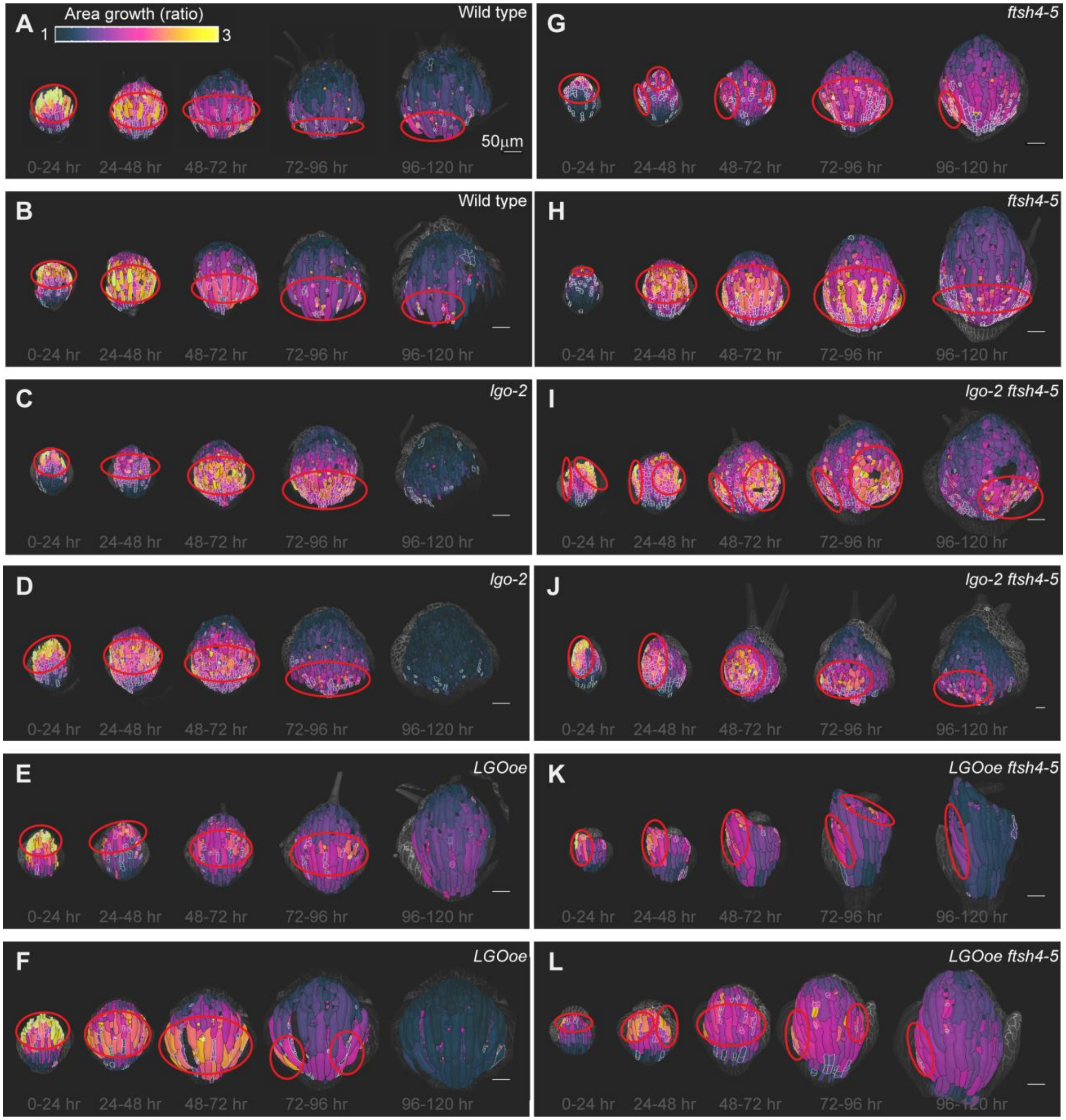


Figure S5: **Cell growth follows a basipetal gradient which is preserved when cell division changes, but altered in *ftsh4-5, lgo-2 ftsh4-5,* and *LGOoe ftsh4-5.*** (A-L) Heat maps of cell area growth over each 24-hour interval that are projected onto the later time point for the remaining two replicates of WT (A-B), *lgo-2* (C-D), *LGOoe* (E-F), *ftsh4-5* (G-H), *lgo-2 ftsh4-5* (I-J), and *LGOoe ftsh4-5* (K-L). The heat map represents the change in ratio of cell area (cell area of later time point divided by cell area of earlier time point) and the scale is 1 to 3. Localization of fast growth is marked by red outlines, and is band-like in WT, *lgo-2*, and *LGOoe* and patchy in *ftsh4-5*, *lgo-2 ftsh4-5*, and *LGOoe ftsh4-5*. The scale bar is 50μm.

Daughter cells that result from a division over a given time interval area outlined in white. (D) Distribution of cell area growth for each time interval. Related to Figure 5.

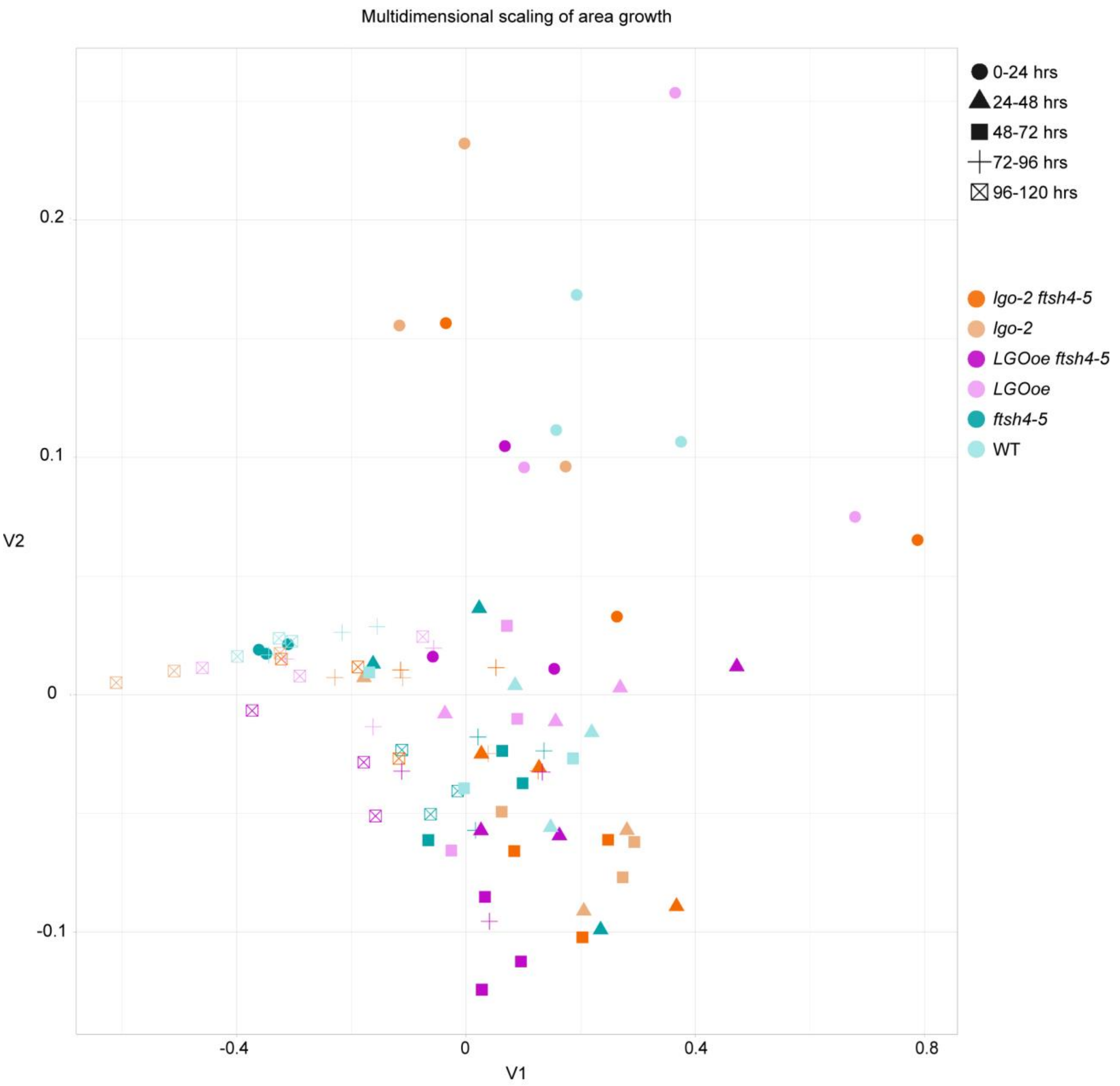


Figure S6: **Distributions of cell growth cluster loosely based on developmental time, and not by genotype.** Multidimensional scaling (also called principal coordinate analysis) is used to represent the distributions of cell area growth rates for each genotype and time interval in two-dimensional space. Increased distance between points indicates larger differences between the growth rate distributions whereas points clustered together indicates that these distributions are similar. Colors and shapes represent genotypes and time intervals, respectively.  Related to Figure 5.

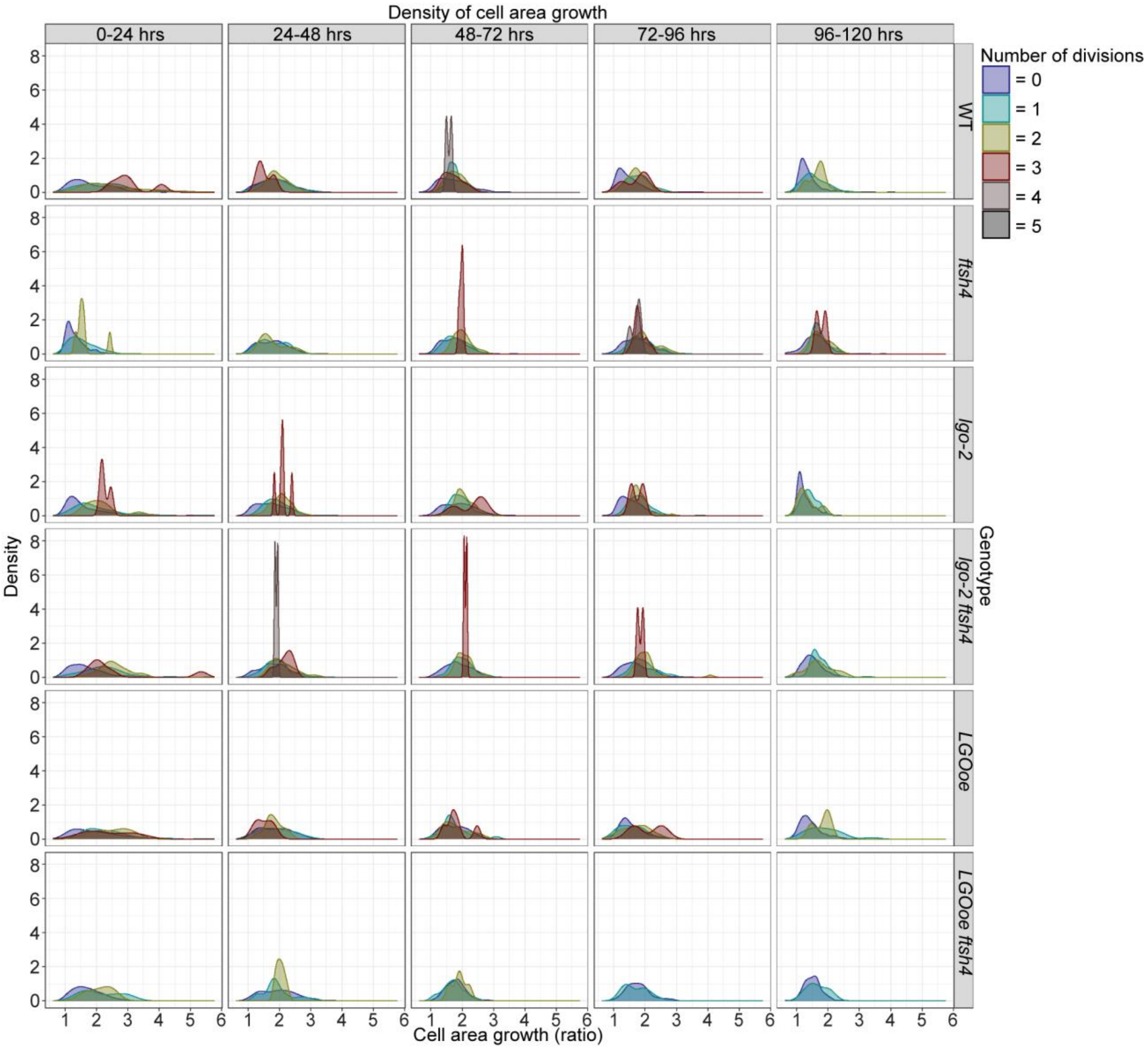


Figure S7: **Cell lineages that are dividing have slightly faster growth rates.** Distribution of cell growth ratio for populations of cells that have divided a specified number of times over a 24-hour interval are plotted for each genotype and time interval. Note that at early time points cell lineages that have undergone more divisions also tend to exhibit higher growth rates, but at later time points the growth curves are more equivalent. Related to Figure 5.

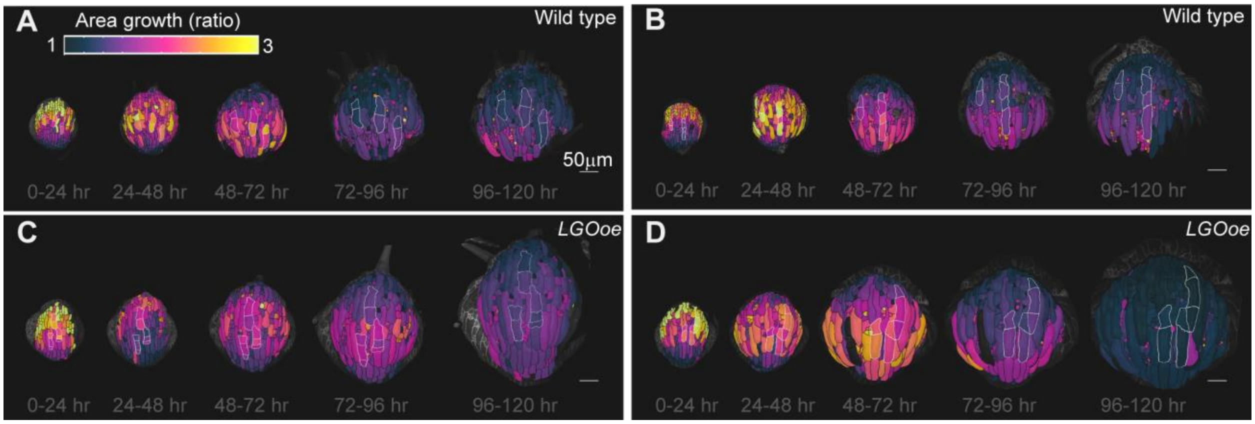


Figure S8: **Cells can have regions with different growth rates in both WT and *LGOoe*.** (A-D) Giant cells are artificially subdivided into multiple cells and outlined in white. The heat maps of cell area growth over each 24-hour interval are projected onto the later time point for remaining two replicates of WT (A-B) and *LGOoe* (C-D). The heat map represents the change in ratio of cell area (cell area of later time point divided by cell area of earlier time point) and the scale is 1 to 3. The scale bar is 50µm. Related to Figure 6.

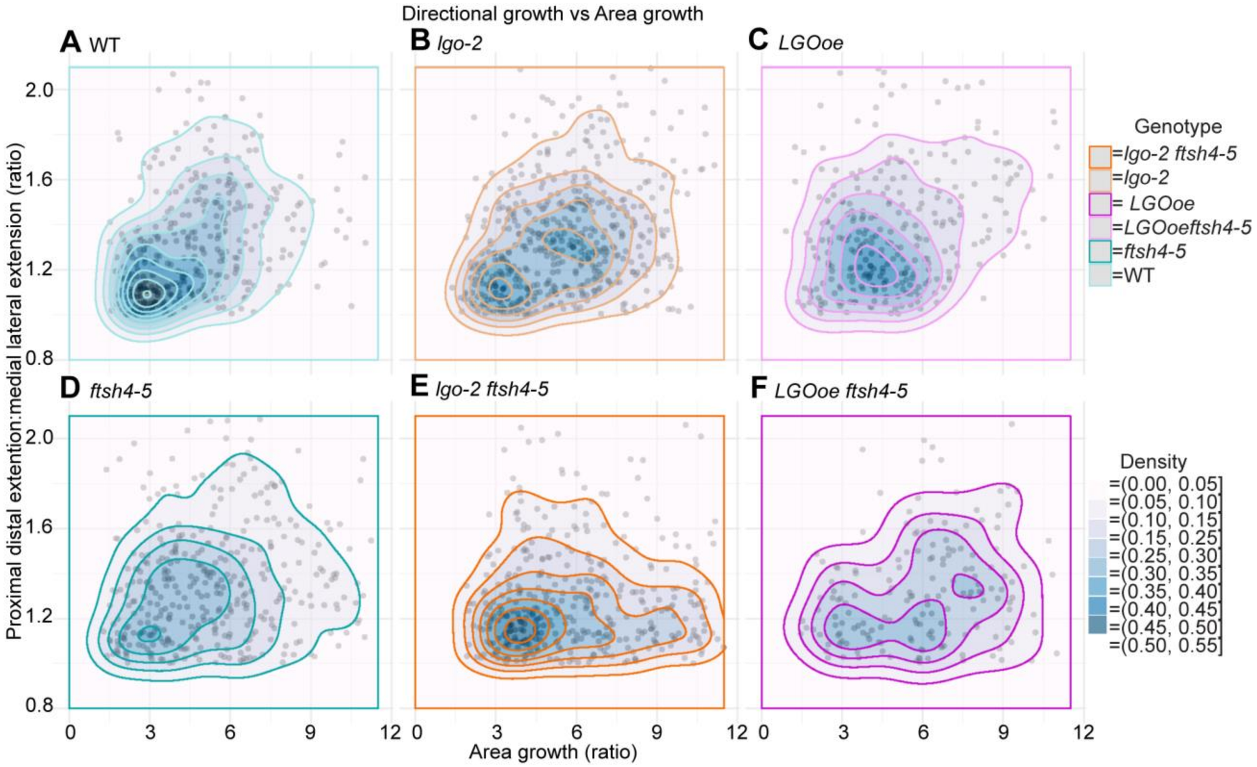


Figure S9: **WT, *LGOoe*, and *lgo-2* have a stronger relationship between growth rate and proximal distal growth direction than *ftsh4-5*, *lgo-2 ftsh4-5*, and *LGOoe ftsh4-5*.** (A-F) Scatter plots of the ratio of proximal-distal to medial-lateral growth (y axis) and the area growth ratio (x axis) for each cell from the 24-hour time point to the 96-hour time point for WT (A), *lgo-2* (B), *LGOoe* (C), *ftsh4-5* (D), *lgo-2 ftsh4-5* (E), and *LGOoe ftsh4-5* (F). Density of the scatter plot is overlaid to show the trends.
